# Fate transitions in *Drosophila* neural lineages: a single cell road map to mature neurons

**DOI:** 10.1101/2021.06.22.449317

**Authors:** Graça S. Marques, José Teles-Reis, Nikolaos Konstantinides, Patrícia H. Brito, Catarina C. F. Homem

## Abstract

Neuron specification and maturation are essential for proper central nervous system development. However, the precise mechanisms that govern neuronal maturation remain poorly understood. Here, we use single-cell RNA sequencing, combined with a conditional genetic strategy to analyse neuronal lineages and their new born neurons in the *Drosophila* larval brain. A focused analysis on the transcriptional alterations that occur right after neuron generation revealed that neuron maturation starts shortly after neuronal birth, with transcription, but no translation, of mature neuronal features such as neurotransmitter (NT) genes. Using NT gene Choline acetyltransferase as an example, we show that the timings of translation initiation are not solely dependent on neuron age but are rather coordinated with the animal developmental stage. This study is the first characterization of the initial phases of neuron maturation, supporting a model where neuron maturation is coordinated with the animal developmental stage through post-transcriptional regulation of terminal effector genes.

## INTRODUCTION

Brain development is a highly complex process that requires the coordinated generation and maturation of thousands of different types of neurons and glia to ensure the formation of complex neuronal circuits. During brain development, a small number of neural stem cells (NSCs) gives rise to the large neuronal diversity found in the adult brain (reviewed in e.g. (Contreras et al., 2019; Holguera and Desplan, 2018). Neural lineages play pivotal roles in neuron fate determination, as transcriptional and molecular changes that occur at each step of lineage progression can be inherited and ultimately determine cell fate specification (reviewed in (Figueres-Oñate et al., 2021; Holguera and Desplan, 2018)). Studies in *Drosophila* have been fundamental to show how the type of neuron generated by each neural lineage is determined by the combination of several layers of transcription factors (TFs) and signaling pathways acting at the level of the NSC, the intermediate progenitors, or even at the level of the differentiating neuron (Baumgardt et al., 2009; Kohwi and Doe, 2013; Landgraf and Thor, 2006; Liu et al., 2019, 2015; Truman et al., 2010; Wang et al., 2014). These core mechanisms of neural lineage progression and neuronal fate specification have been shown to be well conserved in mammals (Bonnefont and Vanderhaeghen, 2021; Briscoe and Novitch, 2008; Cau and Blader, 2009; Kimura et al., 2008; Kohwi and Doe, 2013). After the specified neurons are born, neurons must undergo a maturation period, during which they establish their final neuronal properties such as axonal and dendrictic arborization and find synaptic partners (Allan and Thor, 2015; Hobert, 2016). This maturation requires the coordinated expression of a combination of effector molecules such as cell surface molecules, ion channels and neurotransmitter (NT) receptors to ensure that neuronal wiring occurs only when their synaptic partners are formed (Kratsios et al., 2015; Kurmangaliyev et al., 2020). In *Drosophila* neurons are formed in two waves, the first occurs in embryos to form the CNS of the embryo itself and the larva (primary neurogenesis), while the second wave occurs in larvae and pupa and is responsible for the generation of the majority of the adult neurons (secondary neurogenesis) (Truman and Bate, 1988). Although secondary neurons are formed during a large developmental time window, neurons remain immature until mid-pupal stages when synaptogenesis synchronously starts (Hartenstein, 1993; Kurmangaliyev et al., 2020; Muthukumar et al., 2014; Özel et al., 2015, 2019).

To date, very little is known about the mechanisms driving the early stages of neuronal maturation (Bradke et al., 2020). Interestingly, it has been recently shown in *Drosophila* that the age-related transcriptomic diversity of neurons is partially lost as early as 15h after neuron birth, resulting in a transcriptomic convergence in mature neurons (Özel et al., 2020). This highlights the need to study young neurons as the transcriptomic profiles involved in the initial phases of neuron maturation can be quickly lost and no longer be detectable in adult neurons.

Previous transcriptomic studies have not been designed to analyse the initial phases of neuron maturation, as previously generated bulk-seq and single cell RNA-Seq (scRNA-Seq) datasets of the developing larval brain either analyse both primary (mature) and secondary (immature and maturating) neurons, not allowing for their clear distinction (Avalos et al., 2019; Harzer et al., 2014; Yang et al., 2016), or analyse brains from the adult animal which contain only post-mitotic fully mature neuronal cells (Allen et al., 2020; Davie et al., 2018; Konstantinides et al., 2018). The remaining available scRNA-Seq atlases specifically analysed the optic lobe (OL) (Konstantinides et al., 2018; Kurmangaliyev et al., 2019, 2020; Özel et al., 2020), or focused only on a sub-set of midbrain lineages (Michki et al., 2021);.

To characterize the transcriptional changes in secondary neural lineages we focused on the developing *Drosophila* central brain (CB) and ventral nerve cord (VNC). The CB and VNC generate one third of all CNS neurons , and are responsible for forming brain structures, such as the mushroom bodies and central complex, critical for memory-directed behavior and navigation (Cognigni et al., 2018; Seelig and Jayaraman, 2015).

We devised a conditional genetic strategy to label, select and sequence the transcriptomes of CB and VNC NSCs, called neuroblasts (NBs) in *Drosophila*, and their immediate progeny, including only 0h to 12.5h-old neurons (a time window prior to neuron transcriptomic convergence (Özel et al., 2020)). Using the Chromium system (10x Genomics) we obtained ∼12.6K single-cell transcriptomes of NBs and their daughter cells, ∼7.2K of which were neurons.

The analysis of the young neurons in our dataset has allowed us, for the first time, to transcriptionally characterize the initial phases of neuron maturation. It revealed that shortly after birth young secondary larval neurons start transcribing several terminal effector genes characteristic of mature neurons, such as ion channels and NT genes, even though they are only required at synaptogenesis which starts days later at pupal stages (Chen et al., 2014; Hartenstein, 1993; Muthukumar et al., 2014; Özel et al., 2015, 2019). We further show that NT genes, such as Choline acetyltransferase (ChaT), is transcribed but not translated in young neurons and that translation of ChAT protein does not solely depend on neuron age, but is rather coordinated with the animal developmental stage only starting in pupal stages P6-P7 (∼48h after puparium formation, APF). Based on these results we propose that the initial stages of neuron maturation can be sub-divided into 3 phases: a first phase when newly born neurons do not transcribe terminal neuronal effector genes; a second phase, shortly after birth, when neurons start transcribing, but not translating terminal neuronal effector genes; and a third phase when terminal effector genes, such as ChAT, start being translated in coordination with the animal developmental stage.

## RESULTS

### Early fate atlas of of NB lineages in CB and VNC

To characterize the initial transcriptional changes that drive larval neural lineage differentiation and early maturation of secondary neurons we performed scRNA-Seq. We have specifically labeled, isolated and sequenced the transcriptome of 3^rd^ instar larval CB and VNC NBs, intermediate progenitors and their newly born neurons. To ensure that only the lineages that originate secondary neurons were analyzed, we devised a conditional genetic strategy that was precisely controlled at a spatial and temporal level, as it is described next. We used the CB and VNC NB specific Vienna Tile Gal4 line#201094 (VT#201094; Figure 1A) to drive the expression of CD8::GFP specifically in NBs. As GFP protein is stable for several hours it is inherited by the NB progeny, effectively labeling neural lineages (Figure 1A). To control the time window of GFP expression, we included a temperature sensitive (ts) tubGal80, which allows GFP expression at 25°C, but represses GFP expression at 18°C (McGuire et al., 2003, 2004). With this conditional genetic system, we could precisely control when NBs start expressing GFP and thus start generating GFP-labeled progeny.

**Figure 1.**
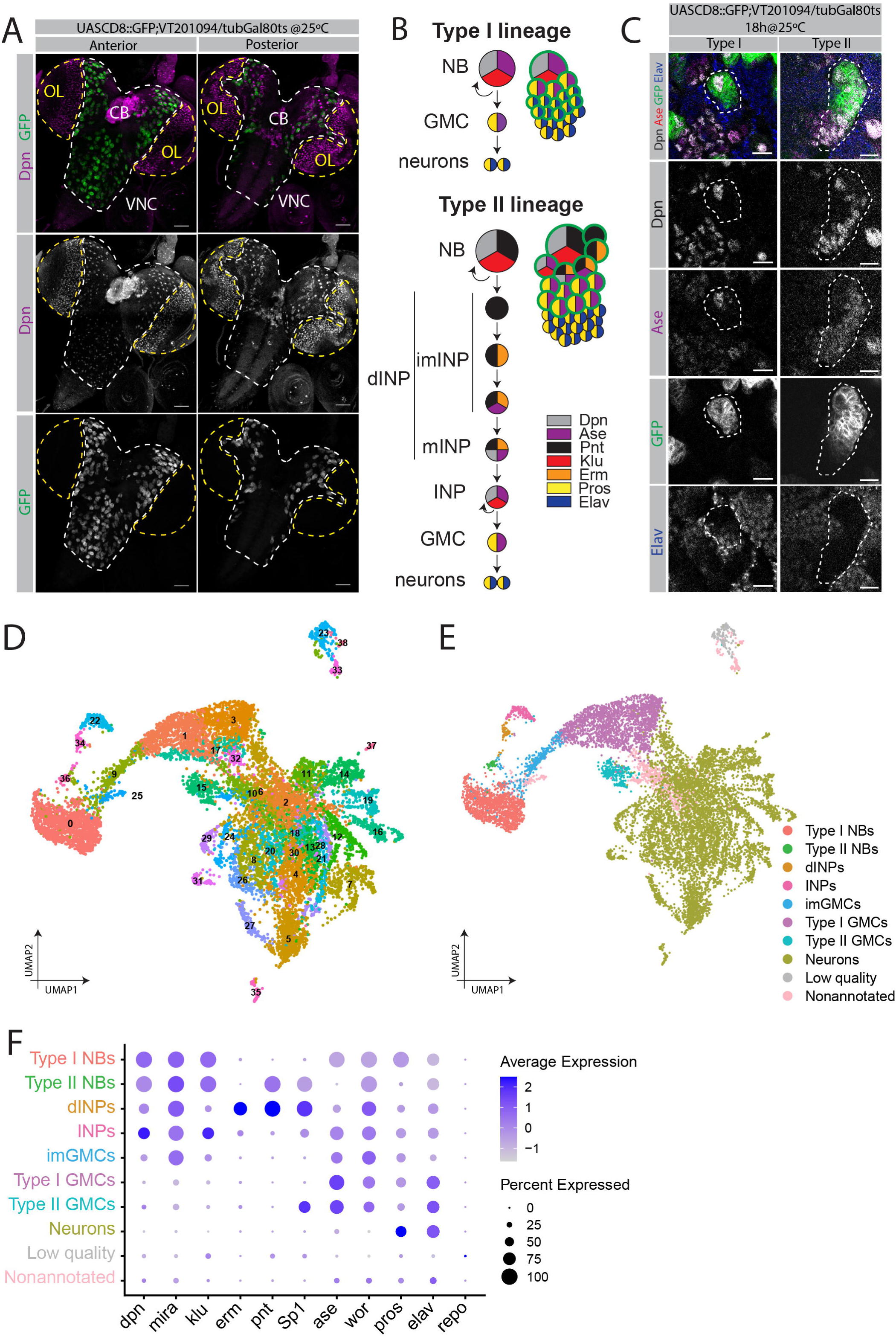
Type I and type II NB lineages identified by scRNA-Seq in wandering larvae brain. (A) Pattern of VT201094-Gal4 driver expression in the whole brain, showing GFP expression in NB lineages from CB/VNC, but not OL. Views from anterior and posterior sides; dpn (purple), GFP (green); scale bar, 50 µm. (B) Schematic representation of type I and type II lineages; cells colored by expression of dpn (grey), ase (purple), pnt (black), klu (red), erm (orange), pros (yellow) and elav (blue); green outline indicates GFP expression and thus the cells in which VT201094 expression is driven in a 18h time window. (C) Close up of type I and type II neural lineages in the anterior side of CB; dpn (white), ase (purple), GFP (green), elav (blue); scale bar, 10 µm. (D) UMAP visualization of final datset composed of 12.7K cells of neural lineages from CB and VNC. The 39 clusters are labeled by number. (E) UMAP plot with clusters grouped based on cell type annotation. Color code as described. (F) Dot plot showing the genes that were used to identify each cell type. CB – central brain, VNC-ventral nerve cord, OL – optic lobe, NB – neuroblast, INPs – intermediate neural progenitors, dINP – developing INPs, GMCs – ganglion mother cells, imGMCs – immature GMCs.

For our experimental set-up, precisely staged wandering 3^rd^ instar larvae were used (equivalent to 105h after larval hatching-ALH at 25°C): animals were raised at 18°C and shifted to 25°C 18h prior to dissection to initiate GFP expression in NBs. Based on the durations of cell divisions, which have been precisely described in the CB (Homem et al., 2013), 18h allow for NBs to divide 9-12 times depending on lineage type, and for several GFP positive neurons to be generated. Neural lineages can be sub-divided mainly in type I, which populate both CB and VNC, and type II lineages, that can only be found in the CB. Type I NBs (approximately 100 NBs per CB lobe and 25 NBs in each thoracic VNC hemisegment (Ito et al., 2013; Truman et al., 2010; Yu et al., 2013) divide every ∼1.3h to self-renew and generate ganglion mother cells (GMCs), which after ∼4.2h divide to form neurons and glia (Homem et al., 2013) (Figure 1B,C). Consequently, in a 18h labeling window, the oldest neurons can be at most ∼12.5h-old (18h-1.3h-4.2h=12.5h), although most likely they are slightly younger as the expression of GFP protein under UAS-Gal4 takes approximately 3h to occur (personal communication, M. Garcez, 03.2021). Type II NBs (8 per CB lobe) divide every ∼1.6h to self-renew and generate an additional intermediate neural progenitor cell (INP); in turn, the INP requires ∼6.6h to mature and divide to self-renew and generate GMCs that will form neurons and glia (Figure 1B,C) (Homem et al., 2013). In type II lineages, due to the extra differentiation cell state (the INPs), the 18h time window allows for the labeling of NBs, INPs and GMCs but fewer neurons (Figure 1B,C). However, the fact that we predominantly do not observe type II GFP labeled neurons after immunofluorescence analysis (Figure 1C) indicates that the majority of neurons represented in this dataset are from type I origin.

We sorted the labeled neural lineages by FACS, based on their size and GFP expression. Two samples were processed in parallel using the Chromium system (10x Genomics) and analyzed with standard Seurat pipeline (Satija et al., 2015; Stuart et al., 2019). Quality control (QC) metrics attested an overall good quality of the samples, such as low percentage of mitochondrial genes (**Figure S1A**) and allowed us to set appropriate filters, resulting in a library of 12,671 cells and 10,250 genes. Using Seurat’s graph-based clustering approach we identified 39 clusters with distinctively expressed marker genes, which were visualized in low dimensional space using the Uniform Manifold Approximation and Projection (UMAP) algorithm (McInnes et al., 2018) (Figure 1D). Moreover, the 2 samples seemed to be evenly distributed through the different clusters, which strengthens the good quality of our dataset (**Figure S1B**). By checking the expression of known marker genes, we were able to identify and annotate the different cells that make up neuronal lineages (Figure 1E,F). Interestingly the two clusters with highest feature number, cluster 0 and 36 (**Figure S1C**) were annotated as NBs. NBs were globally identified by the expression of known NB markers *dpn*, *mira*, *wor* and *klu* (all gene symbols according to Flybase; Figure 1F). Type I NBs were distinguished as they additionally co-express *ase* (cluster 0; Figure 1F, **S1D**), while type II NBs do not (cluster 36; Figure 1F, **S1D**). Type II NBs and their lineage INPs and GMCs, were additionally identified by the expression of *Sp1* and *pnt* (Figure 1F).

In our analysis INPs (type II lineages) were separated into two clusters, which based on their expression patterns we named “developing INPs” (dINPs, cluster 34; Figure 1F, **S1D**), and “INPs” (INPs, cluster 22; Figure 1F, **S1D**). The “dINP cluster” includes several stages of INP maturation including immature INPs expressing *erm* and *ase*, and the more mature INPs that express *erm*, *ase* and also *dpn*. The “INP cluster” includes the fully mature INPs, which stop expressing *pnt* and *erm* and turn on *klu* expression (Berger et al., 2012).

GMCs were identified based on their expression of *wor* and *ase*, but absence of *dpn* and *mira* (Figure 1F). Type II GMCs were additionally identified by their expression of *Sp1* (cluster 15; Figure 1F, **S1D**), in contrast to type I GMCs which do not express *Sp1* (clusters 1, 3, 17, 32; Figure 1F, **S1D**). As GMCs do not have any specific marker, usually being identified by combinatorial expression of several genes, it has been difficult to specifically isolate and analyze them. Our transcriptome that includes both type I and type II GMCs thus represents a good tool for the study of GMCs and lineage transition from self-renewing NBs to terminally dividing GMCs.

Neurons were identified based on their expression of *pros* or *elav*, but absence of *dpn*, *mira* and *wor* (clusters 2, 4-8, 11-14, 16, 18-21, 24, 26-31, 35, 37; Figure 1F, **S1D**). Finally, we identified a group of cells as an intermediate state between type I NBs and GMCs (cluster 9, **Figure S1D**). This annotation is supported by the location of this cluster in the UMAP plot between type I NBs and GMCs (Figure 1D, E). Like type I NBs, they express the NB marker *mira* (∼92%), but only one third still express *dpn* (∼35%), an essential marker of NBs (**Figure S1D**). As these cells also differ from GMCs, where both *dpn* and *mira* are not expressed, this led us to annotate them as immature GMCs (imGMCs; Figure 1E,F). Notably, the presence of this transitional state indicates that the fate commitment of type I GMCs does not occur immediately after asymmetric NB division and is rather a progressive process.

In our analysis we did not identify any cluster enriched for the glial cell marker repo, indicating that no glial cells were included in our dataset (Figure 1F). Neural lineages also generate glial cells (Doe, 2017; Enriquez et al., 2018), thus, their absence might have been due to the limited temporal window analyzed or to the fact that newborn glia might not yet express characteristic markers such as *repo*.

Clusters characterized by the expression of ribosomal subunit genes, which are associated with low quality (Ilicic et al., 2016), were annotated as “low quality”, and clusters with no obvious expression of any of the previously mentioned markers were identified as “nonannotated” (Figure 1E,F).

This transcriptomic single cell atlas represents, to our knowledge, the first specific characterization of 3^rd^ instar CB and VNC neural lineages and their new born secondary neurons.

### Validation and characterization of imGMCs as a transition state between type I NBs and GMCs

Next, we wanted to test if the cluster identified as the transient imGMCs could be identified *in vivo*. Based on their UMAP location, imGMCs are transcriptionally “between” type I NBs and GMCs (Figure 1E). The partial absence of *dpn*, a NB marker, indicates that imGMCs are no longer fully committed to being a stem cell and supports these cells’ transitional state into a more committed cell fate.

The presence of this intermediate cell state was validated by us *in vivo* by immunofluorescence. In some lineages, the daughter cells immediately next to type I NBs are Dpn^+^Mira^+^ (Figure 2A, arrow). We could unambiguously distinguish NBs and their daughter cells, as daughter cells have a smaller size when compared to their progenitor type I NBs. The low number of these double Dpn^+^Mira^+^ imGMCs is in accordance with our transcriptomic data, where these Dpn^+^Mira^+^ imGMCs should only represent ∼33% of imGMCs. This is in line with other studies that reported Dpn being retained in the nucleus of the daughter cells after NB division in type I lineages of the anterior region and attributed to newborn GMCs (Boone and Doe, 2008).

**Figure 2.**
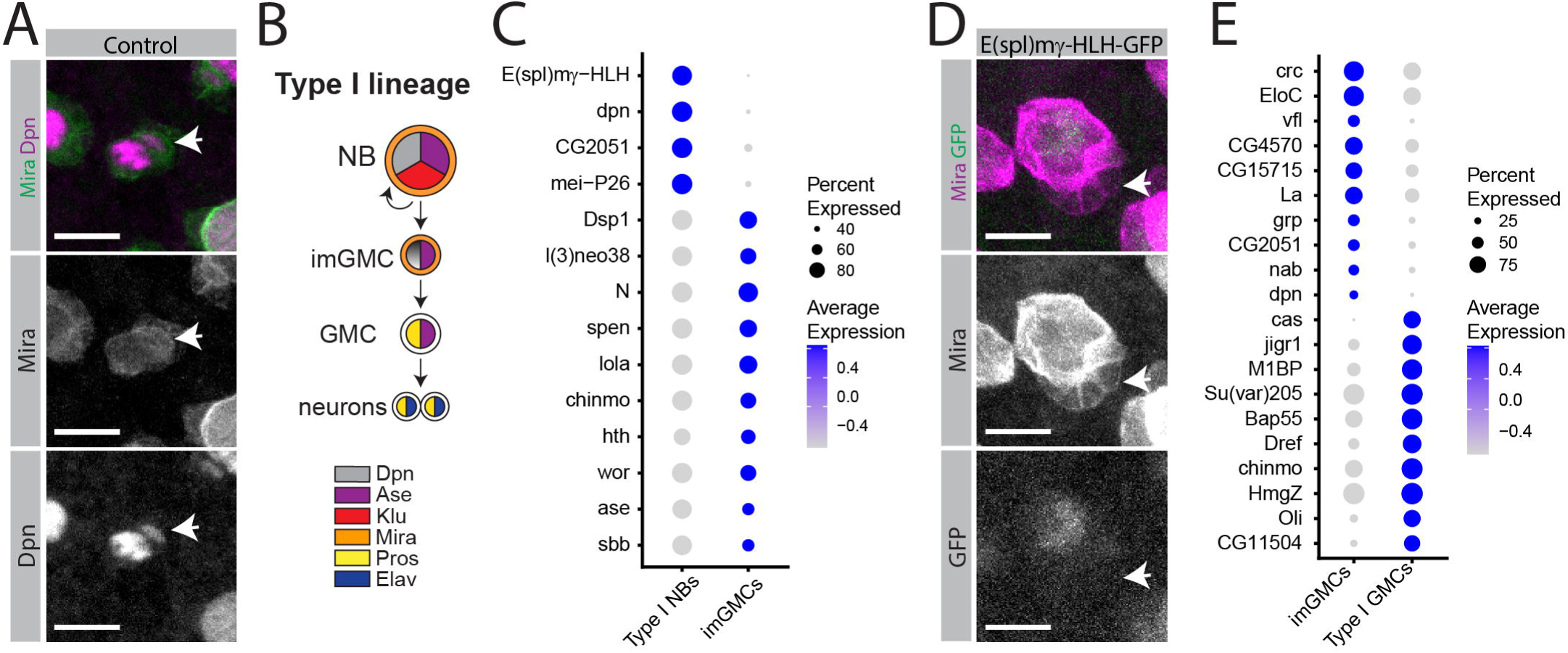
Characterization of NB-imGMCs-GMCs transition in type I lineages. (A) Close up of a type I neural lineage with one imGMC. Type I NBs are Dpn^+^Mira^+^, and upon division, transient imGMCs (white arrow) retain Dpn expression; mira (green), dpn (purple); scale bar, 10 µm. (B) Schematic representation of type I lineages including imGMCs; cells colored by expression of dpn (grey/gradient), ase (purple), klu (red), mira (orange), pros (yellow) and elav (blue). (C) Dotplot showing the top TFs/DNAB differentiating type I NBs *vs.* imGMCs. (D) E(spl)mγ-HLH is expressed in type I NBs, but not in imGMCs (white arrow); mira (purple), E(spl)mγ-HLH reporter (green); scale bar, 10 µm. (E) Dotplot showing the top TFs/DNAB differentiating imGMCs *vs.* type I GMCs. NB – neuroblast, INPs – intermediate neural progenitors, dINPs – developing INPs, GMCs – ganglion mother cells, imGMCs – immature GMCs.

Consequently, these results prove that the bioinformatic analysis of our dataset can efficiently discriminate transient cell states, such as imGMCs (Figure 2B).

The transition from NB to GMC involves dramatic cellular changes, as the loss of self-renewal capacity (Prokop and Technau, 1991). We aimed at identifying genes that might be regulating this transition. Having determined that the transition from NB to GMC is a stepwise process, we compared the transcriptomes of imGMCs to their mother cells, the type I NBs, and to their “downstream” GMCs. We focused our analysis in differentially expressed transcription factors (TFs) and DNA binding genes (DNAB) as several TF/DNAB have been described to be master regulators of neural lineage fate transitions (Eroglu et al., 2014; San-Juán and Baonza, 2011; Weng et al., 2010).

The comparison between imGMCs and type I NBs identified *E(spl)mγ*, a Notch target expressed in NBs (Zacharioudaki et al., 2012), as being downregulated in imGMCs (Figure 2C). To confirm this result, we performed an *in vivo* analysis of a protein reporter for *E(spl)mγ*, which showed that *E(spl)mγ* is expressed in type I NBs, but its expression is no longer seen in the rest of the lineage, not even in the youngest GMCs closest to the NB, which positionally corresponds to the imGMCs (Figure 2D).

On the other hand, imGMCs have increased expression of several co-transcriptional repressors (as *Dsp1*, *l(3)neo38*, *spen* and *sbb*) and genes involved in neuronal differentiation (as *chinmo*, *lola*, *ase*) (Figure 2C). The expression of transcriptional co-repressors is consistent with the transcriptional changes required for the transition to a more committed fate (Rives-Quinto et al., 2020). The comparison between imGMCs and mature GMCs has also identified several differentially expressed genes. As predicted from being a transitional state, imGMCs express higher levels of NB markers as *vfl* (or *zelda*, *zld*), *nab* and *dpn* (Reichardt et al., 2018) then their mature counterparts (Figure 2E). imGMCs also express higher levels of *EloC* (Figure 2E), involved in RNA Polymerase II (Pol-II) elongation, for which knockdown has been shown to cause a reduction in GMC number and shorter lineages (Neumüller et al., 2011). Interestingly, mature GMCs express higher levels of *Su(var)205* (or HP1; Figure 2E) which has been recently shown to be recruited by Pros to promote heterochromatin compaction and terminal neuronal differentiation in GMCs (Liu et al., 2020). Supporting their increased neuronal commitment, GMCs also express higher levels of several neuron differentiation genes than imGMCs (e.g., *jigr1* and *chinmo*; Figure 2E).

Overall, these comparisons suggest that the transition from NBs to GMCs involves a very fast downregulation of Notch, progressive transcriptional and chromatin silencing and heterochromatin formation, with simultaneous upregulation of several neuronal differentiation genes. It further identifies several differentially expressed genes which are candidate regulators of the NB to GMC transition.

### Identification of candidate regulators of neuronal lineage progression by differential gene expression analysis

To identify additional cell specific markers and regulators of neural lineage progression, we performed a comparative analysis of the transcriptomes for all intermediate cell states in both type I and type II lineages. We focused in the most differentially expressed TFs and DNAB genes between cell states, as previously done for imGMCs.

#### I. Type I GMCs vs. neurons

We analysed the transition from GMCs to neurons in type I NB lineages. As expected, GMCs, being less differentiated than neurons, show increased expression of genes associated with neural proliferation, such as *N*, *ase* and *wor* (Figure 3C). HmgD, a chromosomal protein involved in DNA bending and chromatin organization, was previously shown to be differentially expressed between NBs and neurons at the transcriptomic level (Yang et al., 2016). Our analysis confirms that *HmgD* remains highly expressed in GMCs, being only downregulated at the transition from GMCs to terminally differentiated neurons (Figure 3A).

**Figure 3.**
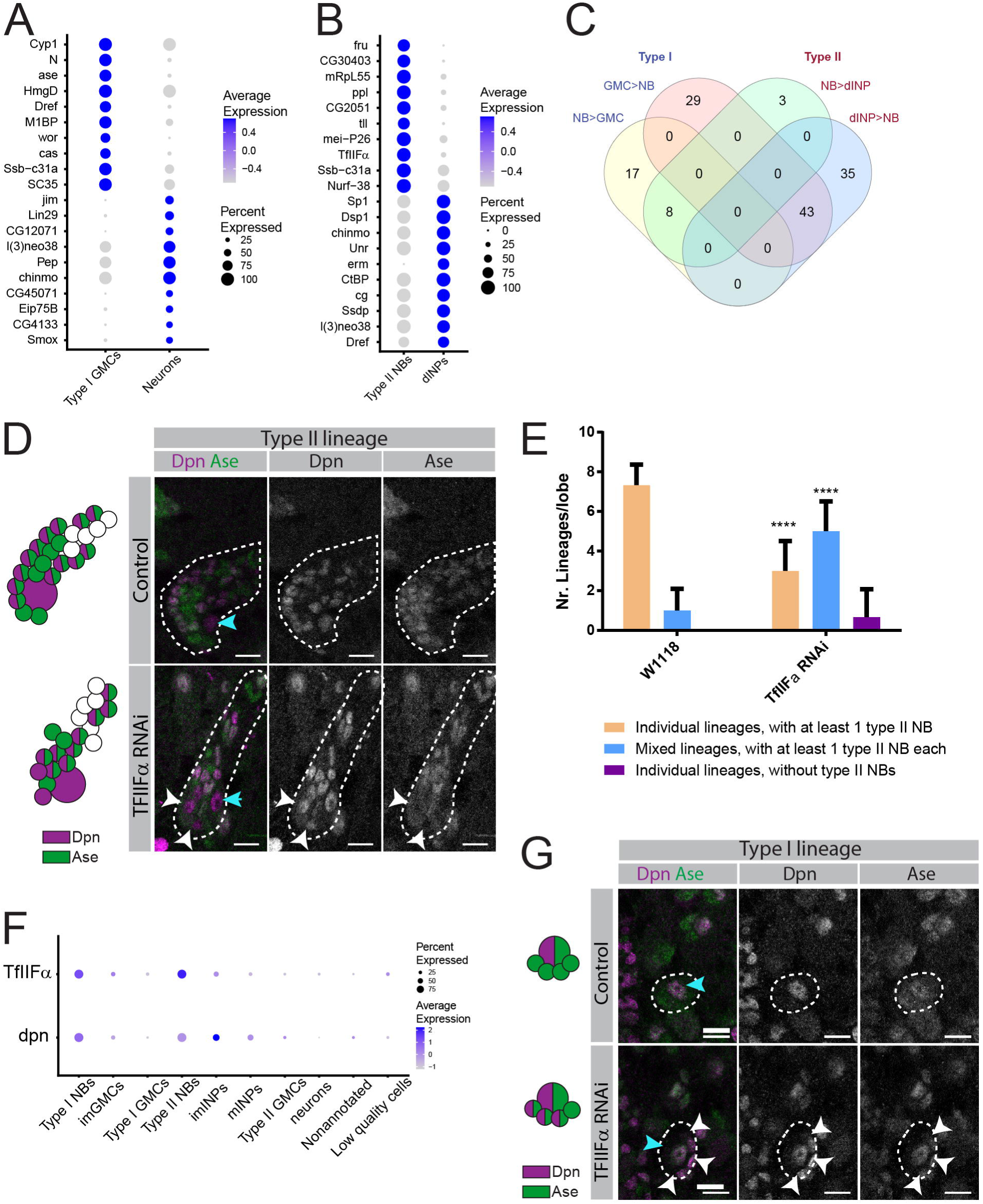
Differentially expressed markers throughout the different differentiation states in NB lineages. (A) Dotplot showing the top TFs/DNAB distinguishing type I GMCs *vs.* neurons. (B) Dotplot showing the top TFs/DNAB distinguishing type II NBs *vs.* dINPs. (C) Venn diagram comparing differentially expressed TFs/DNAB genes, in common between type I NBs *vs.* type I GMCs and type I NBs *vs*. dINPs. (D) Type II lineages of 3^rd^ instar larva; control (upper panels) and TfIIFα RNAi (lower panels); schematic representations (left) and immunofluorescence images (right); Dpn (purple), Ase (green); blue arrowheads indicate primary type II NBs; white arrowheads indicate examples of ectopic NB-like cells; scale bar, 10 µm. (E) Quantification of abnormal morphologies of type II lineages expressing TfIIFα RNAi. Control: average of 6 independent lobes; TfIIFα RNAi: average of 9 lobes from 7 independent brains. Error bars represent ± SD, **** *p* < 0.0001 (two-tailed unpaired *t* test). (F) Dot plot showing the expression of *TfIIFα* throughout the differentiation stages of type I and type II lineages. Dpn expression shown as reference. (G) Type I lineages of 3^rd^ instar larva; control (upper panels) and TfIIFα RNAi (lower panels); schematic representations (left) and immunofluorescence images (right); Dpn (purple), Ase (green); blue arrowheads indicate primary type II NBs; white arrowheads indicate examples of ectopic NB-like cells; Scale bar, 10 µm. NB – neuroblast, INPs – intermediate neural progenitors, dINPs – developing INPs, GMCs – ganglion mother cells, imGMCs – immature GMCs.

Conversely, the top genes that are more highly expressed in neurons *vs.* GMCs, include genes related to neuronal differentiation, such as *Jim*, *Lin29* (or *dati*) and *Smox* (or *Smad2*) (Iyer et al., 2013; Schinaman et al., 2014; Zheng et al., 2006).

#### II. Type II NBs vs. dINPs

Type II lineages are characterized by the presence of INPs, which are able to divide 4-6 times to self-renew and generate GMCs, which in turn originate neurons and glia (Bello et al., 2008; Boone and Doe, 2008; Bowman et al., 2008). Even though several genes have been identified as important regulators of type II lineages (Álvarez and Díaz-Benjumea, 2018; Bayraktar and Doe, 2013; Bayraktar et al., 2010; Eroglu et al., 2014; Hakes and Brand, 2020; Weng et al., 2010; Zhu et al., 2011) and the single cell transcriptome of type II INPs, GMCs and neurons has been described (Michki et al., 2021), a full understanding of what leads to the formation of an INP is lacking.

To characterize the transcriptomic changes that occur at the NB to INP transition we compared type II NBs and dINPs (immature and maturating INPs). This comparison identified several genes known to be specifically upregulated in dINPs, as *Sp1* and *erm* (Figure 3B) (Álvarez and Díaz-Benjumea, 2018; Weng et al., 2010). Conversely, type II NBs were found to express higher levels of *tll* (Figure 3B), as previously described (Hakes and Brand, 2020). However, a large fraction of the genes found to be differentially expressed between type II NBs and dINPs were also found to be differentially expressed between type I NBs vs. GMCs, suggesting these genes are part of a common genetic program for neural differentiation and not type II specific (Figure 3C).

The comparison between type II NBs and dINPs also revealed that a basal RNA Pol-II transcription factor (Pol-II), *TfIIFα*, is more highly expressed in NBs (Figure 3B). There have been recent reports showing how the rate of basal transcription regulation (e.g., elongation, pausing) is important for cell fate (Bai et al., 2010). Our analysis suggests that there is differential expression of the basal *TfIIFα* between NBs and their more committed offspring, potentially indicating that differential composition of RNA Pol-II complex might be an important mechanism to differentially control transcription in cells with different fates in *Drosophila* neural lineages.

To test whether TfIIFα is important for the fate change from type II NBs to dINPs we expressed RNAi against TfIIFα in type II NBs and their lineages and evaluated the impact of this knock down in lineages *in vivo*. Knock down of TfIIFα in type II NBs leads to defective lineages containing ectopic Dpn^+^Ase^-^ NB-like cells (Figure 3D, white arrowheads), a phenotype absent in control brains (Figure 3D). In addition, the knock down of TfIIFα in type II NBs results in disorganized lineages that frequently get mixed with each other at the level of (d)INPs/GMCs (an average of 5 ± 1.5 mixed lineages per lobe *vs*. 1 ± 1.1 mixed lineages in the control; Figure 3E), thus preventing exact quantification of each differentiation state.

Although TfIIFα was not on the top 20 differentially expressed genes between type I NBs and GMCs, it is still more expressed in type I NBs than in GMCs (Figure 3F). To test if TfIIFα has a conserved role in type I lineages, we knocked it down in type I NBs. In control type I lineages there is one type I NB (Dpn^+^Ase^+^), up to one imGMC (Dpn^+^), and several GMCs (Dpn^-^Ase^+^; Figure 3G). RNAi of TfIIFα in type I NBs causes the formation of up to 3 Dpn^+^Ase^+^ type I NB-like cells, although smaller in size than wild-type NBs (Figure 3H, white arrowhead). This phenotype suggests that these ectopic NB-like cells are progeny that retained Dpn expression after NB division. In summary, TfIIFα RNAi leads to defects in type I and II NB lineage differentiation and super-numerary NB-like cells, showing that TfIIFα plays an important role in lineage progression.

### Transcriptomic differences distinguish cells with different degrees of differentiation

The differential gene expression analysis clearly revealed an increasing level of neuronal differentiation from NBs to neurons. Remarkably, the UMAP plot itself (Figure 1E) recapitulates the *in vivo* NB lineage progression order for both type I and type II lineages: NB→ imGMCs→ GMCs→ neurons and NB→ dINPs→ INPs→ GMCs, respectively. We have further validated the order of lineage progression in our dataset by predicting the future state of each cell using the RNA velocity method (La Manno et al., 2018) (**Figure S2A, B**).

As in the CB and VNC NB division does not occur in perfect synchrony, some of the transcriptional changes that occur along differentiation might be overshadowed. To further identify the gene expression trajectories each cell must go from NB to a mature neuron, we have ordered cells in pseudotime using Monocle 2. This method takes advantage of individual cell asynchronous progression and uses gene expression changes to order cells along a certain trajectory such as lineage differentiation (Trapnell et al., 2014). For this analysis we assessed the clusters that were part of either type I or type II lineages. Both lineages were processed independently and, for both cases, cells were consistently ordered from the less differentiated cells (NBs) to the more differentiated ones (neurons on type I lineages and GMCs on type II lineages, Figure 4A,B**, S2C**). The type II lineage trajectory identified two clear GMC fate branches (Figure 4B). However, manual analysis of the genes differentially expressed between both branches revealed that the same genes are expressed with only slightly different expression levels. We hypothesized that these 2 branches might be an artifact caused by the low number of cells from type II lineages available and to the lack of more differentiated cells (neurons) in the trajectory to allow a more complete comparison.

**Figure 4.**
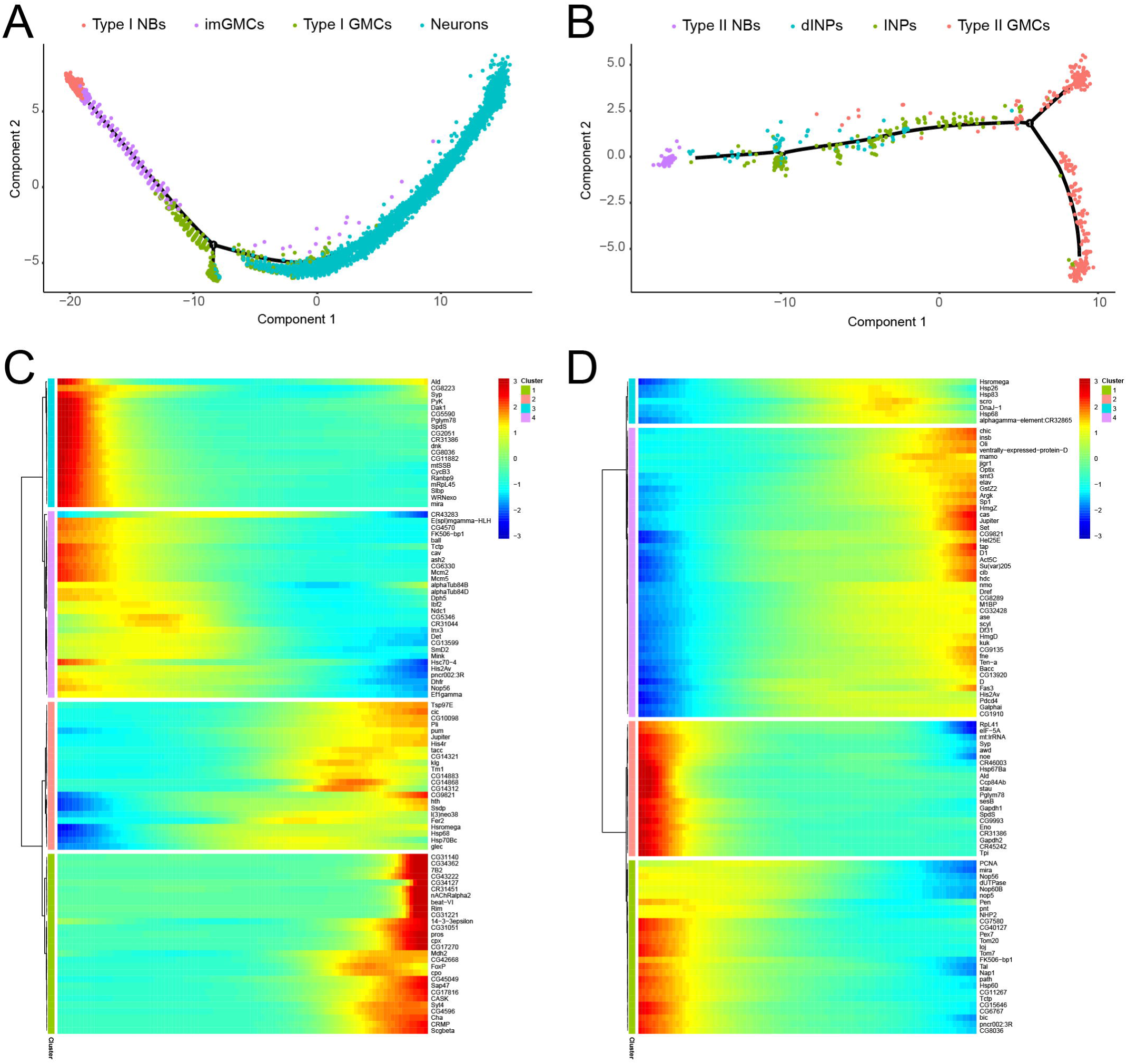
Transcriptomic dynamics of differentiation in type I and type II neural lineages. (A and B) Cell trajectory for type I neural lineages (A) and type II neural lineages (B); cell ordering and trajectory defined based on pseudotime analysis using monocle; colored by cell type. (C and D) Heatmap with the top 100 genes most differentially expressed throughout pseudotime for type I neural lineages (C) and type II neural lineages (D); differential expression analysis performed with monocle. NB – neuroblast, INPs – intermediate neural progenitors, dINPs – developing INPs, GMCs – ganglion mother cells, imGMCs – immature GMCs.

To identify genes that are dynamically expressed along differentiation, we have analysed the top 100 genes that varied the most throughout pseudotime and displayed them in a heatmap. As expected, for both lineage types, known NB fate regulators such as *mira* and *Syp* are more expressed in the cells at the beginning of the trajectory (less differentiated) in comparison to cells at the end of the trajectory (more differentiated; Figure 4C,D). On the other hand, neuron regulatory genes, such as *pros*, are more expressed in more differentiated cells (Figure 4C). Interestingly the analysis of the genes that varied the most in pseudotime in type II lineages identified a small group of genes that seemed to be more expressed prior to GMC generation (Figure 4D, 1^st^ group) and could potentially be important for the INP to GMC fate transition, or potentially represent a transitioning state equivalent to the previously described type I imGMCs. Most of these genes code for heat shock response related proteins (Hsromega, Hsp26, Hsp83, Hsp68 and DnaJ-1). Although the heat shock response was initially thought to be solely dependent on stress conditions, recent evidence suggests that it is cell specific and responds to the proliferative and metabolic needs of the cell (Li and Tennessen, 2017; Li et al., 2016; Ma et al., 2015), which could explain a specific upregulation of heat shock proteins in this major differentiation state transition.

Interestingly, both the RNA velocity method and Monocle suggest that within the neuronal population it is possible to identify different ages or degrees of differentiation. For instance, only the most differentiated neurons of this dataset (Figure 4C; end of trajectory) have an upregulation of genes involved in more specialized roles such as synaptic function and transmembrane transport, as is the case of *CASK*, *cpx* and *nAChRalpha2* (Buhl et al., 2013; Schulz et al., 2000; Sun et al., 2009). Since all neurons in this dataset are younger than 12.5h, these results show that in this short time window it is possible to identify age-related differences between these cells.

### Transcriptomic differences distinguish neurons by age

The transcriptional differences observed among the neuron population in our dataset, suggest that despite their young age, these neurons have begun the process of maturation. A simple analysis to identify the expression pattern of known neuronal markers characteristic of very young neurons, as *Hey*, showed that only the neurons closest to GMCs express this marker (Figure 5A). Hey is a target of Notch, previously shown to be expressed transiently only in Notch^ON^ early born neurons (Monastirioti et al., 2010), and consistently, only approximately half of these very young neurons express *Hey* (Figure 5A). In neuron clusters farther away from GMCs in the UMAP, several genes related to more mature neurons are expressed, consistent with these clusters representing older neurons. This includes several genes coding for terminal neuronal effector proteins as ion transporters and NT pathway members (e.g. *nrv3* and *nSyb*, respectively; Figure 5B, B’). The neuronal clusters furthest from GMCs, as clusters 5 and 35, predicted to be the older neurons in this population, express the highest number of ion channels (Figure 5C**, S3A**) and adhesion molecules (Figure 5C’), essential for axonal development and neuronal circuit formation (Ranscht, 2000; Tan et al., 2015). Furthermore, genes involved in NT activity and biosynthesis, such as the fast-acting NTs *VGlut*, *VAChT*, *Gad1* and *Vmat* are only expressed in clusters farther away from GMCs (Figure 5D). In our neuron population 10,47% express *VGlut*, 8.72% express *VAChT*, 3.15% express *Gad1*, and 0.87% express *Vmat*, markers for glutamatergic, cholinergic, GABAergic, and monoaminergic neurons, respectively (**Figure S3B**). Interestingly these young neurons express only one NT identity, as only a very low percentage of the neurons in our dataset presented counts for simultaneous expression of NT identity marker genes. The assessment of multiple combinations of the 4 fast acting NT revealed that the highest percentage amounted to 0.65% for the simultaneous presence of VGlut and VAChT (**Figure S3B**).

**Figure 5.**
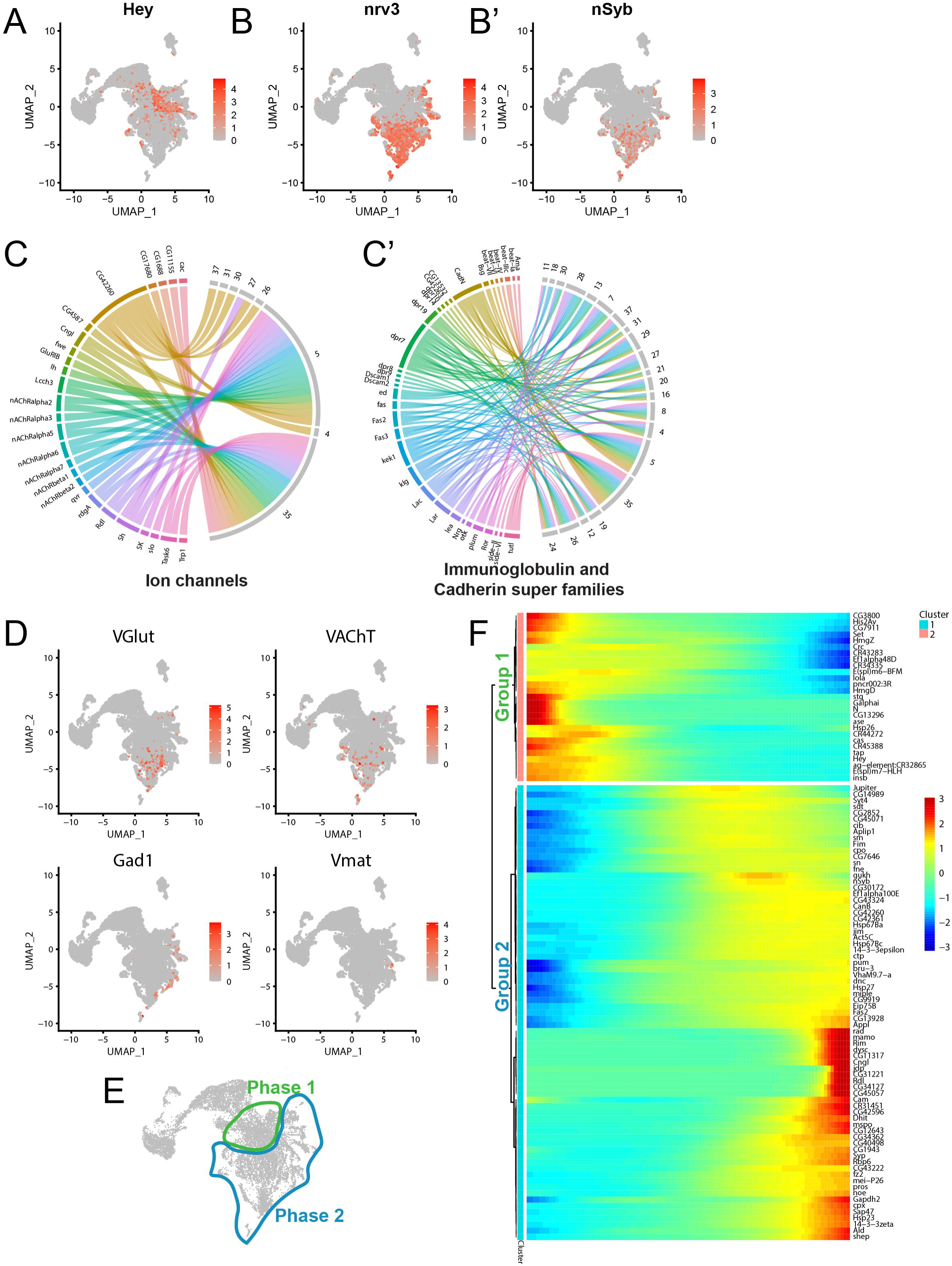
Transcriptomic differences in neurons identify 2 phases of maturation. (A) Feature plot for the young neuronal marker *Hey*. (B,B’) Feature plots for the neuron maturation markers *nrv3* (B) and *nSyb* (B’). (C,C’) Chord diagrams showing the correspondence between Ion channels (C) and Immunoglobulin and Cadherin super families (C’) and the clusters in which they are differentially expressed. (D) Feature plots for neurotransmitter identity markers (*VGlut*, *VAChT*, *Gad1*, *Vmat*). (E) Schematic representation of a UMAP plot showing the division of neurons into the proposed Phase 1 and Phase 2 of neuronal maturation. (F) Heatmap with the top 100 genes most differentially expressed throughout pseudotime in neurons; differential expression analysis performed with monocle.

Overall, the analysis of this neuron population seems to separate them into two sub-groups that for simplicity we have classified into: “Phase 1” of maturation, which includes the very young immature neurons not transcribing terminal neuronal effector genes as NT and ion channels; “Phase 2” of maturation, which includes the older neurons transcribing terminal neuronal effector genes (Figure 5E).

To verify if there is indeed a (clear) transcriptional difference between genes expressed in younger/less mature neurons and older/more mature ones, we have subset the neuronal population and analyzed it independently with Monocle. We identified 2 major tendencies of transcriptional profiles that vary throughout pseudotime, which recapitulate the 2 previously identified maturation phases (Figure 5F). Group 1 represents a transcriptional profile composed by genes that are more highly expressed in cells at the beginning of the trajectory – phase 1 neurons (Figure 5F). A GO analysis showed an enrichment of terms associated with cell fate determination, regulation of neurogenesis and neuron fate commitment (**Table S1**). Group 2 refers to genes that are more expressed in cells at the end of the trajectory, meaning the oldest and more mature – phase 2 neurons (Figure 5F). Group 2 is enriched for terms related with synaptic transmission and NT regulation (**Table S2**). Overall, the transcriptional analysis of young secondary neurons revealed that neurons begin their maturation process shortly after birth, a process marked by the expression of NT-associated genes and several other terminal neuronal effector genes.

### Neurons in phase 2 of maturation transcribe the neurotransmitter gene ChAT, but do not translate it into protein

The expression of NT genes, ion channels and other terminal neuronal effector genes in secondary neurons in larval stages was surprising as these neurons will only initiate synaptogenesis days later in pupal stages (Muthukumar et al., 2014). We sought to determine if the presence of mRNA in these young phase 2 neurons is already accompanied by the expression of the respective protein. We used the fast-acting neurotransmitter choline acetyltransferase (ChAT) as a case-study, as its mRNA is rarely present in phase 1 neurons and is expressed in phase 2 neurons (Figure 6A). As ChAT is a well-established marker of cholinergic neurons (Hamid et al., 2019; Salvaterra and McCaman, 1985), this sub-set of neurons likely corresponds to cholinergic neurons, which together with glutamatergic neurons, correspond to the largest class of adult neurons. In addition, a commercial antibody for this protein is available, allowing for the study of its localization *in vivo*.

**Figure 6.**
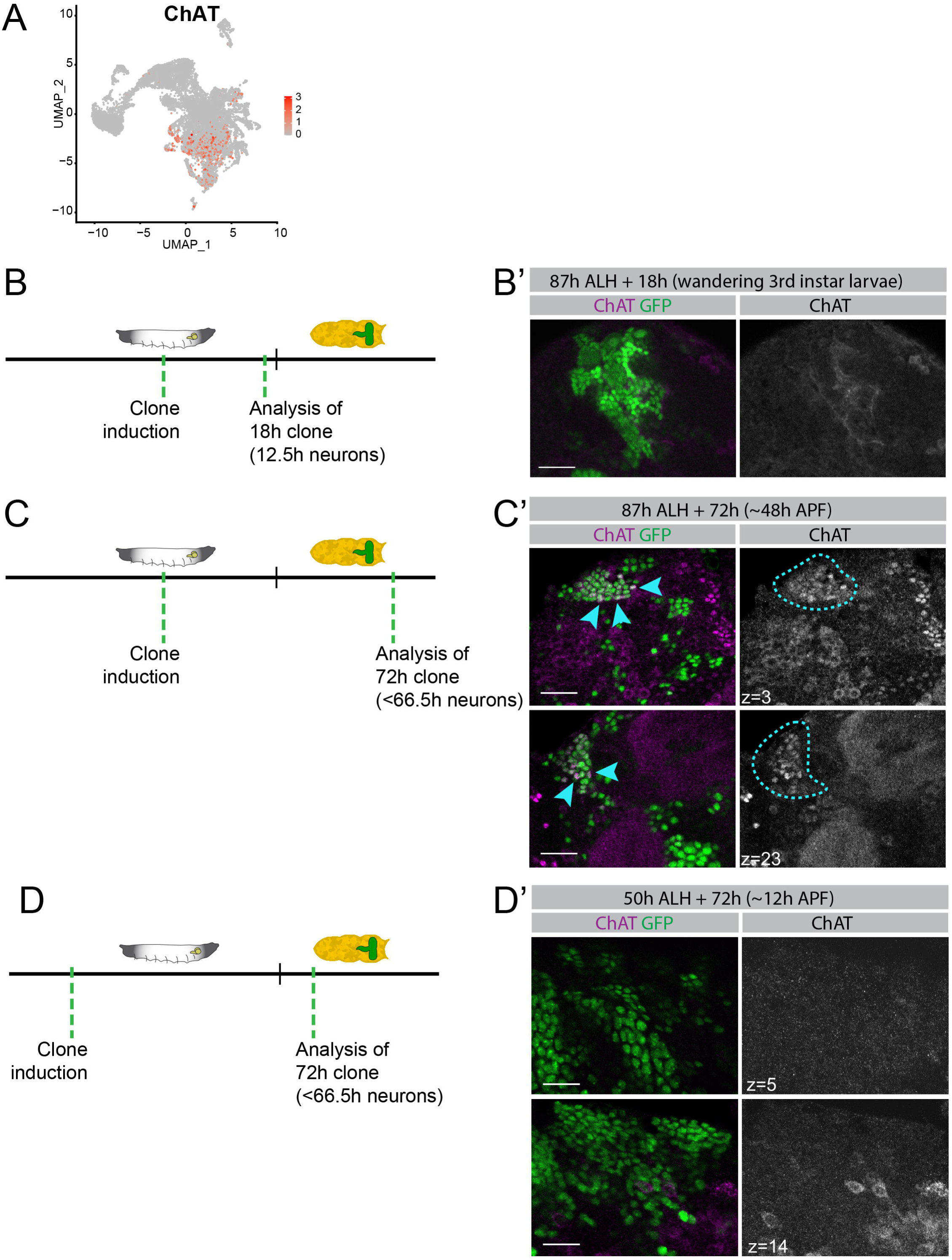
Timing determinants of ChAT translation initiation in neurons. (A) Feature plot for *ChAT*. (B-D’) Clones induced in neural lineages. Schematic representations (B,C,D) and immunofluorescence images (B’,C’,D’). ChAT antibody staining (purple) and GFP (green). (B, B’) 18h-clone induced at 87h and analysed at 105h after larval hatching-ALH (wandering 3^rd^ instar larvae). (C, C’) 72h-clone induced at 87h and analysed at 159h ALH (approximately 48h after puparium formation-APF); two Z-slices of same stack displayed; blue arrowheads indicate examples of GFP positive cells co-expressing ChAT; outline in ChAT pannel outlines clone. (D, D’) 72h-clone induced at 50h and analysed at 122h ALH (approximately 12h APF); two Z-slices of same stack displayed. Scale bar, 20 µm.

We labeled cells using a similar strategy as to generate this atlas, raising the animals at 18°C and shifting them to 25°C 18h before dissection, at the equivalent to 105h ALH (Figure 6B). In this set of experiments, we used a permanent labeling strategy to ensure that all generated neurons in the multiple time windows analyzed remained GFP positive. Interestingly, none of the GFP^+^ neurons generated in the 18h window are also stained with the ChAT antibody, indicating that the protein is not expressed (Figure 6B’). These results indicate that even though the oldest neurons in our dataset (phase 2) already have ChAT mRNA molecules, these are not yet being translated into protein.

### Initiation of ChAT translation is coordinated with animal developmental stage rather than neuron age

The presence of ChAT mRNA but the absence of protein suggests that there is a delay in the initiation of its translation in maturing neurons. To understand what determines translation initiation of ChAT in maturing neurons we tested the following two hypotheses: 1) translation of ChAT is dependent of neuron age and requires neurons to be older/of a certain age; 2) translation of ChAT is coordinated with animal developmental stage starting only when synaptogenesis begins in pupal stages. In order to test these hypotheses, we generated permanent labeling clones which allowed us to fluorescently label neural lineages, and thus neurons from birth, and follow them until a certain age and/or a specific developmental stage. As in 18h clones in wandering 3^rd^ instar larvae there is no expression of ChAT protein in GFP positive neurons (Figure 6B’), we next allowed these neurons to age for a longer time. In order to do this, we induced clone formation at 87h ALH as previously, but instead allowed them to develop for 72h, meaning that the older neurons will be up to 66.5h old, and the animal will be 159h ALH (∼48h APF; Figure 6C). In these clones, co-expression of GFP and ChAT proteins is detected in several neurons (Figure 6C’, arrowheads and outline). However, at this point, these results still fit both hypotheses, as neurons may be expressing ChAT because they are older than in the previous clones, or simply because the animal itself is older and closer to the onset of synaptogenesis (Muthukumar et al., 2014). In order to uncouple neuronal age from the animal’s age, we generated a clone for the same duration of 72h but starting at an earlier time in animal development. We have thus induced clone formation at 50h ALH (Figure 6D). This new timeline still allows neurons to age up to 66.5h-old, but the animal itself, though still a pupa, would be only ∼12h APF (122h ALH). Interestingly, in these clones there is no co-expression of GFP and ChAT (Figure 6D’). These results allowed us to conclude that ChAT protein translation does not start solely depending on the neuronal age itself, being rather dependent on the age of the animal. As we have designed this experiment to allow the neurons in these clones to undergo the larva to pupa transition, which is driven by a pulse of the steroid hormone ecdysone, these results further allow us to conclude that neither the larva to pupa transition, nor the ecdysone pupariation peak, are sufficient to initiate ChAT translation. To further narrow down when ChAT translation initiates, we have also generated a 48h clone at 87h ALH and analysed it at ∼24h APF (**Figure S4A, A’**). Neurons in these clones have very little co-expression of GFP and ChAT.

Based on these data, we propose that within neuronal maturation, there is a “phase 1” where newly-born neurons do not transcribe NTs and other terminal neuronal effector genes, followed by a “phase 2” when neurons start transcribing NT, ion channels and other terminal effector genes. In “phase 2”, and using ChAT as an example, we show that its mRNA is however kept untranslated. Afterwards, approximately at 48h APF, a “phase 3” of maturation starts, when the neuron begins translation of ChAT in a coordinated manner with animal and brain development (Figure 7A). At this point, we can only suggest this model for neuroactive molecules, more specifically for ChAT, but we propose this to be a conserved mechanism for NT genes and possibly other molecules associated with more differentiated neuronal characteristics, such as ion channels. Interestingly several RNA binding proteins and translation regulators such as Syncrip (Syp), musashi (msi), pumilio (pum), brain tumor (Brat), or polyA-binding protein interacting protein 2 (Paip2), are expressed in phase 1 and phase 2 neurons (Figure 7B), which might provide a molecular mechanism for translation inhibition of NT genes during larval and young pupal stages. Although the pupariation pulse of Ecdysone does not seem to be the trigger of the molecular mechanisms that will ultimately drive NT translation, other important pulses of Ecdysone occur during pupal development which might, together with other players, provide for the overarching temporal cue for translation initiation and phase 3 of neuron maturation.

**Figure 7.**
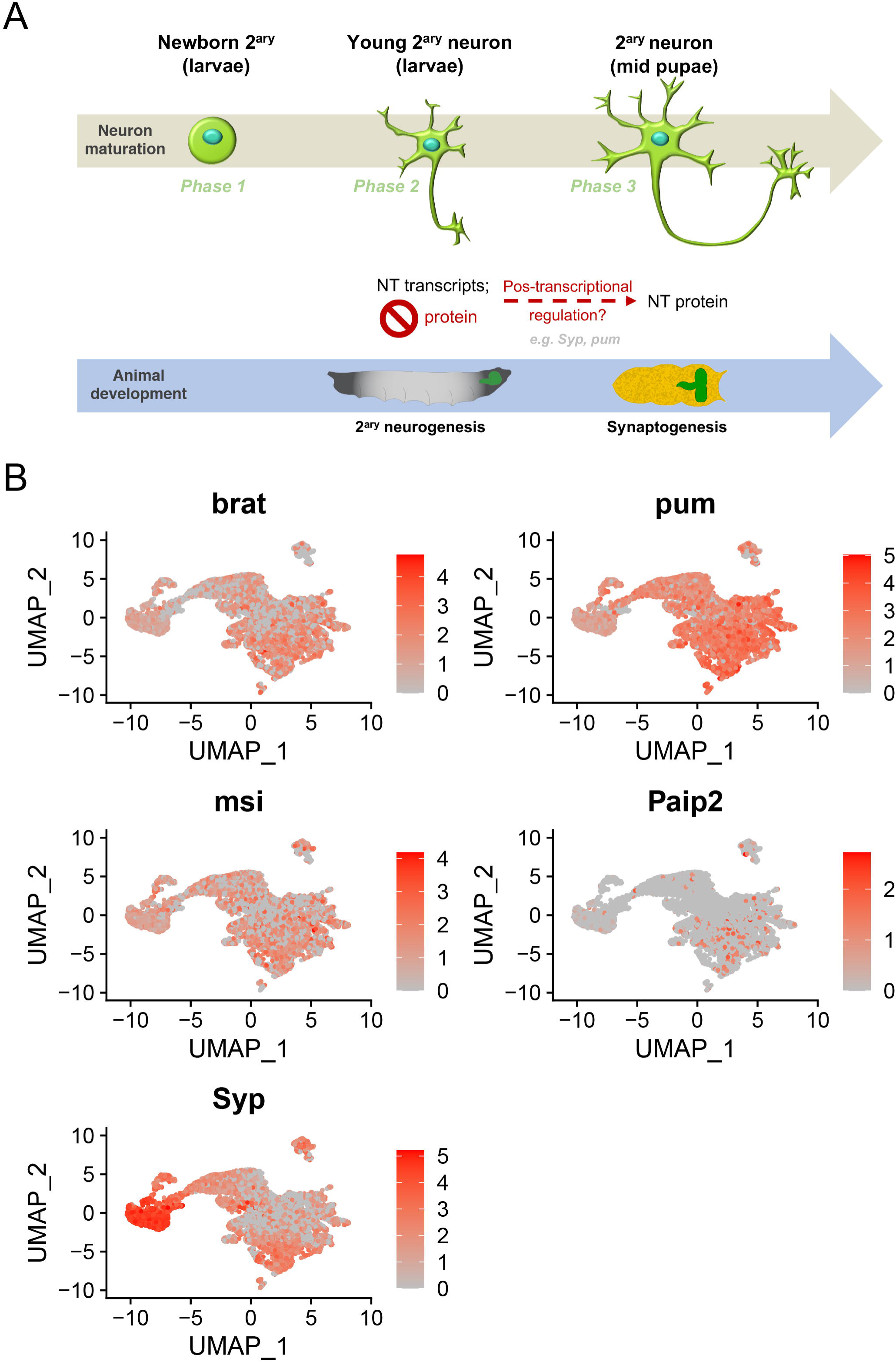
Phases of neuronal maturation in the developing CB and VNC. (A) Model for the initial phases of neuronal maturation. (B) Feature plots to visualize the expression of translation inhibitors *brat*, *pum*, *msi*, *Paip2* and *Syp*.

Altogether these findings reveal that neuron maturation starts very quickly after neuron formation, and sub-divide this process into 3 distinct phases based on the timings for transcription or translation initiation of terminal effector genes. It raises the possibility that neuron maturation is coordinated with the stage of brain development and synaptogenesis by temporal regulation of translation of NT genes and other neuronal terminal effector genes. Such mechanism would ensure that neurons are only fully mature, when their synaptic partners are as well, thus relaying on non-automomous signals to reach full maturation. Transcription without translation may provide an efficient responsive mechanism to ensure that protein synthesis can quickly and coordinately initiate for the final phase of neuron maturation and synapse formation.

## DISCUSSION

In this study we generated a comprehensive atlas of CB and VNC NB lineages in the developing *Drosophila* wandering larval brain. This allowed us to analyze the transcriptional changes throughout neural lineage progression, from the less differentiated NB to young neurons. Moreover, the fact that our temporally regulated labeling strategy allowed us to collect and analyze the neural lineages generated within a short and defined temporal window (18h maximum) means that we were able to greatly remove the contribution of temporal variation from our dataset and analyse precisely aged cells.

Although previous single cell studies of specific aspects of the *Drosophila* brain have been made (Allen et al., 2020; Avalos et al., 2019; Croset et al., 2018; Davie et al., 2018; Konstantinides et al., 2018; Kurmangaliyev et al., 2020; Michki et al., 2021; Özel et al., 2020), our work is the first atlas of CB and VNC neural lineages of the developing brain specifically including only secondary neurogenesis. This atlas represents a valuable resource for studying the regulatory genes and networks involved in NSC proliferation, lineage differentiation, neural cellular diversity and neuron maturation.

Notably, we were able to identify not only previously described cell states within neural lineages, but also a transient cell state, the imGMCs, showing that our dataset has enough resolution to resolve brief states of differentiation. Cases of newborn GMCs retaining Dpn after NB division have already been reported by (Boone and Doe, 2008) in the CB. Somewhat similar occurrences of a NB diving and originating Dpn^+^ cells, not necessarily NBs, have also been described in the OL (Mora et al., 2018; Pinto-Teixeira et al., 2018). Although this transition state between type I NBs and GMCs has already been seen *in vivo*, this is the first study where such population is characterized regarding its transcriptome. Moreover, the presence of this transitional state shows that commitment to a GMC fate does not occur immediately after NB asymmetric division; thus, a time lag after the asymmetric inheritance of basal polarity proteins by the GMC is required for the full establishment of GMCs. Although we did not identify imGMCs in type II lineages, we do not exclude that they may also exist and could have been missed due to the overall lower cell numbers for type II lineages. Since in type II lineages, INPs go through a maturation period themselves, it would not be surprising that type II GMCs would also go through such maturating steps, and it would be interesting to address this in future studies.

By comparing the transcriptional profile of different cell states throughout differentiation we were able to identify possible candidates to regulate lineage differentiation. One example is TfIIFα, a member of the RNA Pol-II machinery, that was identified in our dataset as being differentially expressed in type II NBs *vs.* dINPs and type I NBs *vs.* GMCs, which we have shown to be important to maintain proper larval neural lineage development *in vivo*. TfIIFα has important roles during transcription initiation, elongation and even pausing (Gong et al., 1993; Landick, 2006; Price et al., 1989; Rossignol et al., 1999). Based on its role in different steps of transcription, TfIIFα, either alone or associated with other TFs, might influence transcription start site selection (Freire-Picos et al., 2005; Khaperskyy et al., 2008) or prevente RNA Pol-II pausing (Price et al., 1989; Rossignol et al., 1999). Alterations in the usual rates by which these processes happen, could affect the transcription of genes responsible for NB differentiation, thus justifying the important role of TfIIFα in regulating lineage progression. In the future it will be interesting to study the mechanisms that directly or indirectly link TfIIFα to neural lineage regulation. We have additionally identified several other differentially expressed genes within neural lineages and it would be interesting to test their potential role in lineage progression. Ultimately, such studies will contribute to better understand not only NB proliferation, but also progression towards differentiation.

One of the most interesting findings in this study was determining that young neurons (<12.5h-old) have transcriptional profiles that reflect their different ages and consequently different degrees of maturation. Our analysis led us to propose the sub-division of neuron maturation into 3 phases: a “phase 1”, composed by immature neurons that have yet to start expressing mRNA of more mature neuronal features; followed by a “phase 2” of maturation, composed by the oldest neurons in our dataset, which start expressing mRNA of maturation markers such as genes involved in axonal development, neurotransmitter activity and ion channels; and finally, a “phase 3” of neuronal maturation where not only mRNA, but also the protein of terminal neuronal effector genes is expressed. Using the NT gene ChAT as a case study, we found that although this gene is expressed in “phase 2” neurons in our transcriptome dataset, its protein is not. We have shown that regardless of neuronal age, it is not until approximately 48h APF that ChAT protein starts being expressed (“phase 3”). This indicates that ChAT translation depends on the age of the animal, rather than on the age of the neuron itself. Ultimately, such timeline seems to match the beginning of synaptogenesis, which is described to start around 60h APF in the CB (Muthukumar et al., 2014). The early expression of mRNA of neuron maturation markers was unexpected since the neurons formed in larval stages, meant for the adult brain, only terminally differentiate in the pupal period (Dumstrei et al., 2003; Pereanu and Hartenstein, 2006; Truman, 1992). So, if ChAT protein is only expressed days after, why does its mRNA starts being expressed so much sooner? And how is its translation being inhibited, and how is it then initiated? One hypothesis is that translation inhibitors might be acting during phases 1 and 2 of neuronal maturation, specifically keeping terminal neuronal effector genes untranslated and consequently keeping the neuron in a state that is not yet fully mature, until the time comes when their synaptic partners are formed and ready to connect. This hypothesis is supported by the fact that we can find several RNA biding molecules and translation inhibitors such as *Syp*, *pum* and *brat* expressed in the neurons in our dataset. Interestingly some of these genes, as *pum* and *brat* are known to be involved in the maternal-to-zygotic transition which is also dependent on post-transcriptional regulation and translation inhibition (Murata and Wharton, 1995; Sonoda and Wharton, 2001). Moreover, according to this hypothesis, in order to allow the translation of ChAT and other NT-associated molecules to be possible in “phase 3”, these translation inhibitors would then need to be downregulated or inactivated at the appropriate pupal stage, a model which would need to be confirmed *in vivo* in the future. In such a model an upstream coordinator of this switch in post-transcriptional regulation would be required. Such role could be taken up, for instance, by hormones, which as systemic signals are ideally placed to coordinate animal and organ development. Although we have shown that the pupariation pulse of the hormone ecdysone is not sufficient to initiate ChAT translation, we do not exclude that other pupal pulses of ecdysone may provide such a function, alone or in combination with other signals. Overall, a mechanism of temporal regulation of translation may provide an efficient responsive mechanism to ensure that protein synthesis can quickly initiate for the final phase of neuron maturation and synapse formation. Ultimately, the identification of these 3 different phases of neuronal maturation represents an important foundation for further studies to understand the mechanism and timelines that regulate neuronal maturation.

## METHODS

### KEY RESOURCES TABLE

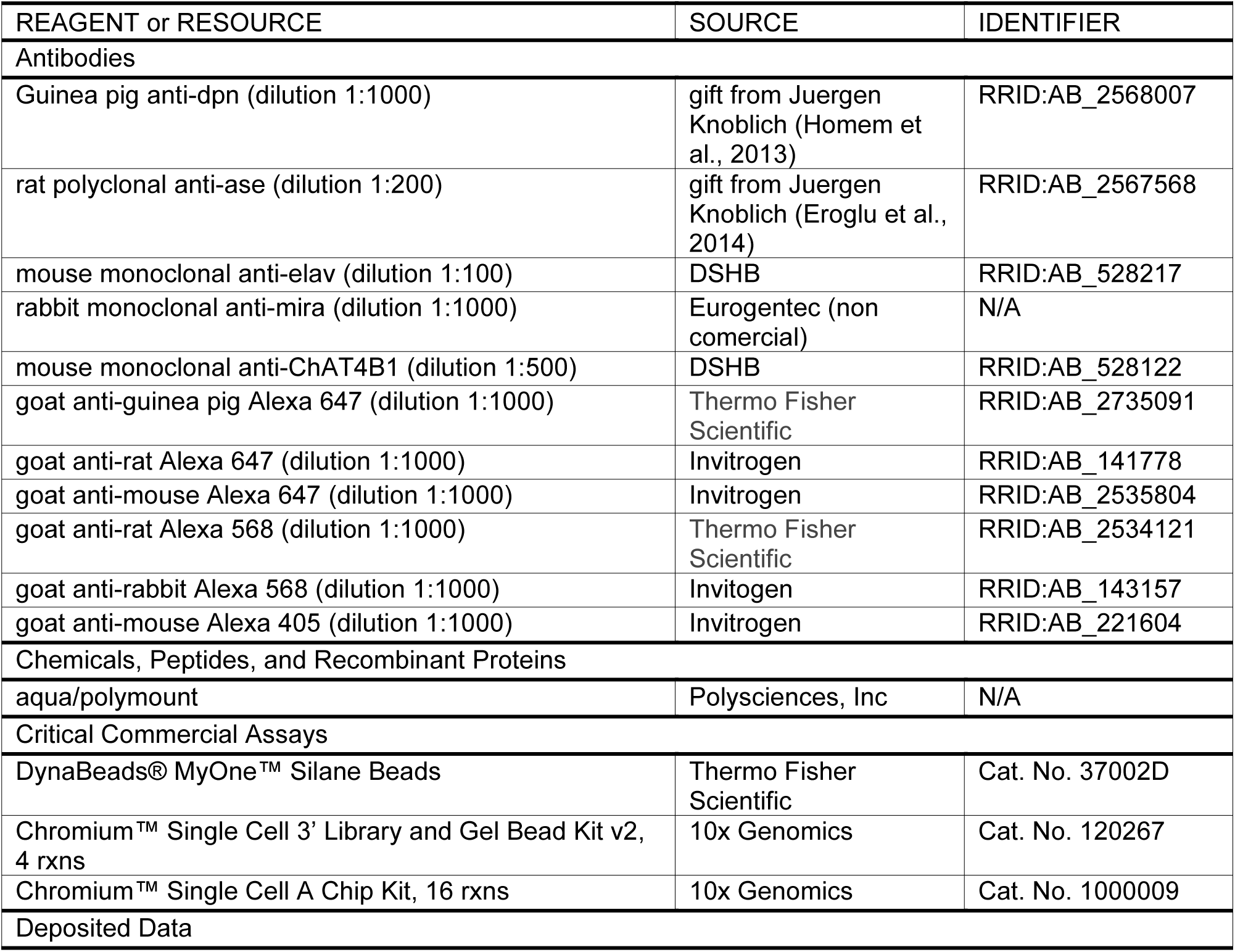
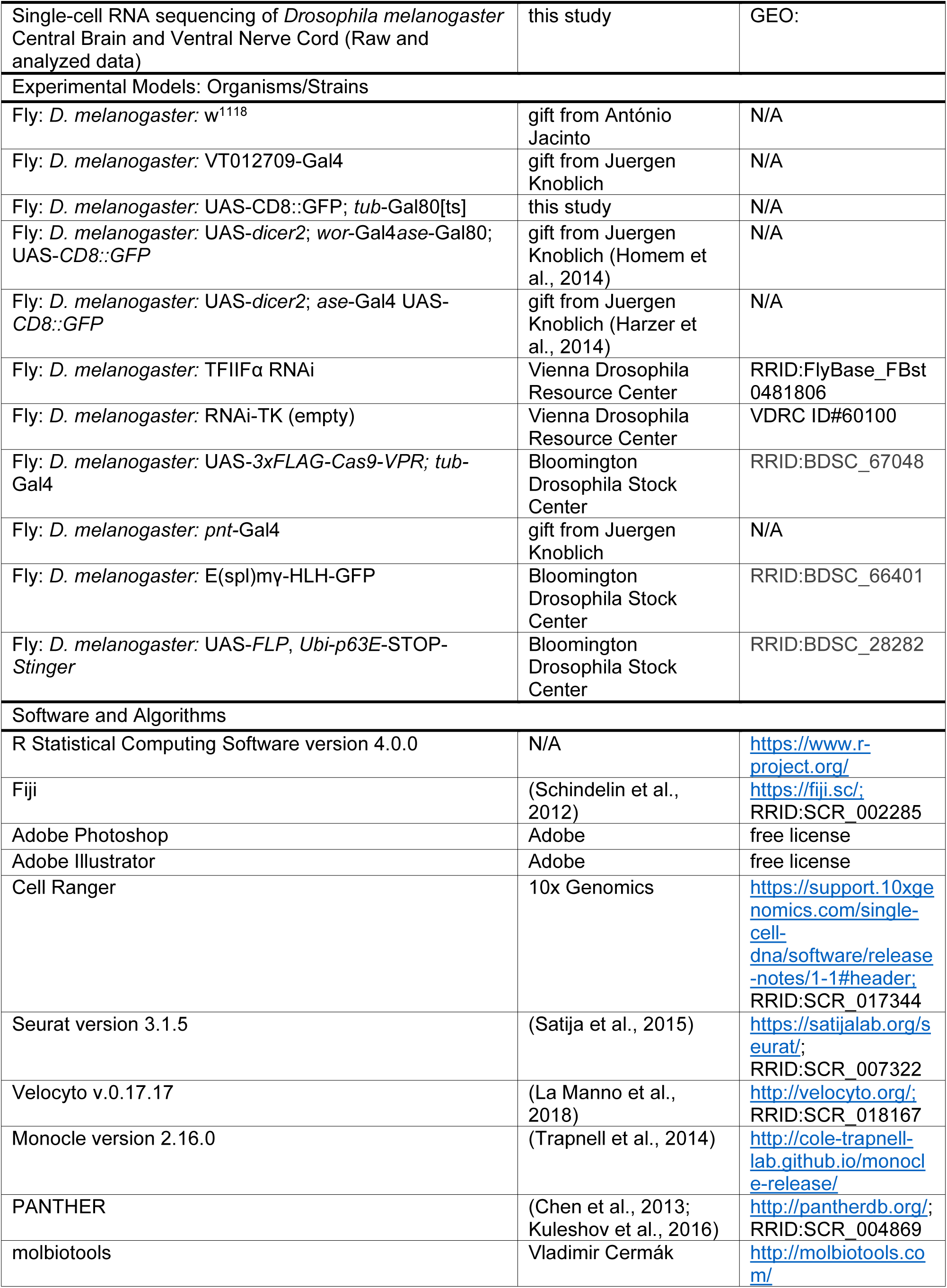
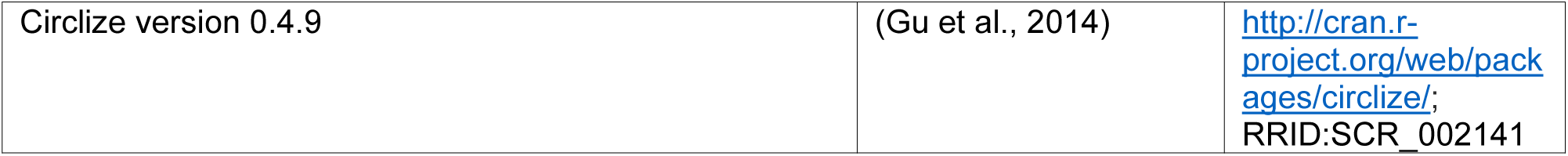

## RESOURCE AVAILABILITY

### Lead Contact

Further information and requests for resources and reagents should be directed to and will be fulfilled by the Lead Contact, Catarina Homem.

### Materials Availability

This study did not generate new unique reagents.

### Data and Code Availability

The datasets generated during this study are available in GEO.

## EXPERIMENTAL MODEL AND SUBJECT DETAILS

### Fly strains

For the scRNA-Seq experiments, temporal labeling of NBs and their lineages was achieved by crossing VT012709-Gal4 (enhancer of Koi and CG15236; CB/VNC NB driver) with UAS-CD8::GFP; tub-Gal80ts males. Fly crosses were set up at 25°C, allowing ∼12h-egg lays. First instar larvae (L1) were synchronized upon hatching and transferred to 18°C to inactivate CD8::GFP expression. CD8::GFP expression was activated 18h previously to the dissection timepoint at 105h after larva hatching (ALH). For the permanent labeling experiments, males from ;;UAS-*FLP*, *Ubi-p63E*-STOP-*Stinger* driver were crossed with ;tub-Gal80ts; VT201094 females and raised as previously described. CD8::GFP expression was activated at 50h or 87h ALH and clones were allowed to form for 18h, 48h or 72h at 25°C.

*w*^1118^ was used as control for RNAi experiments. UAS-*dicer2* ; *ase*-Gal4 UAS-*CD8::GFP* was used as a type I NB driver (Harzer et al., 2014) and UAS-*dicer2* ; *wor*-Gal4 *ase*-Gal80 ; UAS-*CD8::GFP* was used as a type II NB driver (Homem et al., 2014). The RNAi lines were crossed to either the control or type I and type II drivers; fly crosses for RNAi experiments were set up at 29°C to increase UAS-Gal4 expression and fluorescence intensity.

For other experiments the stock line #60100 from VDRC TK collection (empty RNAi vector) and ;UAS*-3xFLAG-Cas9-VPR; pnt-*Gal4 were crossed and used as control. Please check the Key Resources Table for more complete information about the *Drosophila* lines used in this study.

## METHOD DETAILS

### Brain Dissociation and Cell Sorting

One hundred and twenty-four third instar larvae (105h ALH) were collected and dissected in supplemented Schneider’s medium (10% fetal bovine serum (Sigma), 20 mM Glutamine (Sigma), 0.04 mg/mL L-Glutathione (Sigma), 0.02 mg/mL Insulin (Sigma) Schneider’s medium (Sigma)). After dissection, brains were transferred to Chan & Gehring solution (Chan and Gehring, 1971) 2 % FBS, and washed once. After this, they were enzymatically dissociated in Chan & Gehring solution 2 % FBS with 1 mg/mL Papain (Sigma) and 1 mg/mL Collagenase I (Sigma) for 1hr at 30°C. Afterwards, brains were washed once with Chan & Gehring 2 % FBS solution and once more with supplemented Schneider’s medium. After these washing steps, brains were resuspended in PBS (phosphate buffered saline) 0.1% BSA (Sigma) and mechanically disrupted using a pipette tip. The cell suspension was filtered through a 30 μL mesh into a 5 mL FACS tube (BD Falcon) and immediately sorted by fluorescence activated cell sorting (FACS) (FACS Aria II, BD). GFP positive NBs and their lineage were collected in a drop of PBS 0.1% BSA. Since NBs represent a lower percentage of sorted cells when compared to neurons, they were sorted separately in order to assure an enrichment of less differentiated cells in the final pool. Cells were resuspended in 0.1% BSA at a final concentration of approximately 400 cell/μL and immediately processed according to the 10x Genomics protocol.

### 10x Genomics experimental procedure

Approximately 25k of the sorted cells (NB lineages) were used to construct single-cell libraries; libraries were obtained using Chromium Single Cell 3’ reagent Kits v2 (10x Genomics) standard protocol. Cells were equally divided into 2 samples (duplicates) and loaded in 2 wells of a Single Cell A Chip, aiming for an estimated target cell recovery of ∼7k cells. Cells were then partitioned into nanoliter-scale Gel Bead-In-EMulsions (GEMs) and reverse-transcribed using an Eppendorf Mastercycler pro Thermal Cycler (Eppendorf), set for 53 °C during 45 min, 85 °C for 5 minutes and hold at 4 °C. Post reverse transcription incubation GEMs were then broken and the cDNA was recovered and cleaned using Silane DynaBeads (Thermo Fisher Scientific). The next step consisted in amplifying the cDNA, by incubating the samples in a Thermal Cycler programmed for 98 °C during 3 minutes, 10 cycles of 98 °C for 15 sec, 67 °C for 20 sec and 72 °C for 1 min, followed by 72 °C for 1 min and hold at 4 °C. The amplified cDNA was then cleaned using SPRIselect and quantified using a TapeStation (Agilent Technologies). The amplified cDNA was fragmented, end-repared and A-tailed by incubating in a Thermal Cycler at 32 °C for 5 min, 65 °C for 30 min and hold at 4 °C; next, the cDNA went through a double-sided size selection using SPRIselect. Subsequently, the samples went through adaptor ligation, by incubating in a Thermal Cycler at 20 °C for 15 minutes, after which there was a new SPRIselect cleanup step. Afterwards, samples were attributed independent indexes and amplified by PCR using a Thermal Cycler set for 98 °C for 45 sec, 14 cycles at 98 ° for 20 sec, 54 °C for 30 sec and 72 °C for 20 sec, followed by 72 °C for 1 min and hold at 4 °C. The generated library went through a new double-sided size selection by SPRIselect and run on a TapeStation for quality control and quantification.

Both samples were subjected to paired-end sequencing using the NovaSeq 6000 system (Genome Technology Center at NYU Langone Health).

### scRNA-Seq Raw Datasets

Each sequenced sample was processed with Cell Ranger Version 3.0.1 for alignment, barcode assignment and UMI counting. Samples were mapped to BDGP6 reference genome from Ensembl.

### scRNA-Seq Dataset Pre-processing

The filtered gene matrices obtained after Cell Ranger processing were analyzed with R package Seurat 3.1.5. Only cells that had at least 200 unique feature counts were included in the analysis. Moreover, we only kept cells with a percentage of mitochondrial genes inferior to 20%. Higher percentages of mitochondrial genes are usually indicative of cell damage/rupture, and consequently, of altered overall transcriptional content (Ilicic et al., 2016). These initial quality control steps resulted in a dataset with 12,671 cells and 10,250 genes.

Samples were normalized using the NormalizeData function, and the top 2000 most variable features were then identified using the FindVariableFeatures function. Next, ScaleData was used to scale all genes; within this step, the percentage of mitochondrial genes was also regressed out, in order to avoid artefacts in subsequent analysis.

### scRNA-Seq Dataset Clustering

We performed a Principal Component Analysis (PCA), using the previously calculated top 2000 most variable genes. Next, we used an elbow plot and a jackstraw approach to identify significant PCs. For the complete dataset analysis, we used the first 51 PCs, as not to include PC that was not significant (p > 0.05). Within the FindClusters function, the resolution parameter was set to 1.55 as it resulted in a granularity that allowed the identification of smaller cell populations such as type II NBs and INPs. The combination of these parameters originated 39 clusters.

Clusters were annotated into major groups corresponding to the different cell types identified in our dataset, resulting in 10 major clusters. This annotation was performed based on well described markers for each cell type (Figure 1F), as well as based on relative cell localization within the UMAP plot.

### RNA Velocity Dynamics

RNA velocity analysis was performed with the python version of Velocyto v.0.17.17 package (La Manno et al., 2018). We used the subcommand ‘velocity run’ to create a loom file for the cells that survived the filtering steps of Seurat pipeline using the *Drosophila melanogaster* genome annotation file (Drosophila_melanogaster.BDGP6.88.gtf) and the bam file with sorted reads that was estimated using the default parameters of the Cellranger software (10x Genomics). We masked repetitive regions using the genome expressed repetitive annotation file downloaded from UCSC genome browser. The loom file created separates molecule counts into ‘spliced’, ‘unspliced’ or ‘ambiguous’. To estimate RNA velocity parameters, we adapted the pipeline used in the analysis of the mouse hippocampus dataset from La Manno et al., 2018. We started by removing cells with extremely low unspliced detection requiring the sum of unspliced molecules per cell to be above the 0.4 threshold. We also selected genes that are expressed above a threshold of total number of molecules in any of the clusters requiring 40 minimum expressed (spliced) counts in at least 30 cells, after which we kept the top 3000 highly expressed and variant genes on the basis of a coefficient of variation CV vs mean fit that uses a nonparametric fit (Support Vector Regression). We applied a final filter to the dataset by selecting genes on the basis of their detection levels and cluster-wise expression threshold. This filter kept genes with unspliced molecule counts above a detection threshold of 25 minimum expressed counts detected over 20 cells, and with average counts of unspliced and spliced expression bigger than 0.01 and 0.08 respectively in at least one of the clusters. Finally, both spliced and unspliced counts were normalize for the cell size by dividing by the total number of molecules in each cell, and multiplying the mean number of molecules across all cells to. All filtering steps resulted in a dataset of 12604 cells and 1086 genes to be used in the RNA velocyto analysis. For the preparation of the gamma fit we smooth the data using a kNN neighbors pooling approach (velocyto subcommand knn_imputation) and k=500 with calculations performed in the reduced PCA space defined by the top 99 principal components. Velocity calculation and extrapolation to future states of the cells was performed under the assumption of constant velocity. Analysis pipeline can be obtained from the corresponding author.

### Single-cell Trajectories

Pseudotime analysis was performed using R package Monocle v2.16.0. The Seurat object, containing all filtering and clustering information was imported to Monocle and subset accordingly. For the analysis of type I lineage the subset included clusters annotated as “Type I NBs”, “imGMCs”, “Type I GMCs” and all clusters identified as “Neurons”. For the analysis of type II lineage, the subset included “Type II NBs”, “dINPs”, “INPs” and “Type II GMCs”; no neuronal clusters were included as, the duration of our labeling combined with the longer cell division timings within type II lineage (Homem et al., 2013), likely did not allow any neurons resulting from type II lineages to be labeled. For the analysis of the neuronal population, the subset included all clusters annotated as “Neurons”, and within those, only cells with 0 counts for *wor* and *repo* were processed.

The 3 subsets (type I lineage, type II lineage and neurons) were then processed independently, but using the same pipeline. Differences in mRNA across cells were normalized and “dispersion” values were calculated using the functions estimateSizeFactors and estimateDispersions, respectively.

To construct single cell trajectories, we started by using the differentialGeneTest function to extract the genes distinguishing different clusters; these genes were then marked to be used for clustering in subsequent calls by using setOrderFilter. Afterwards, dimensions were reduced by using the Discriminative Dimension Reduction Tree (DDRTree) method, and finally, ordered using the orderCells function. In order to perform differential expression analysis, we used differentialGeneTest again, however, this time to obtain the differentially expressed genes as a function of pseudotime. These genes were then ordered by qvalue, and the top 100 hits were presented for each type of lineage.

### Gene annotation lists

TFs/DNAB genes were selected from the Gene List Annotation for *Drosophila* (GLAD) (Hu et al., 2015), obtained from //www.flyrnai.org/tools/glad/web/. The list for ion channels was also obtained from GLAD.

For the cadherin super family analysis, we used a gene list from FlyBase, obtained under the group “CADHERINS” – FBgg0000105.

For the immunoglobulin super family analysis, genes were selected according to FlyBase FBrf0167517 (Vogel et al., 2003).

In all three cases, gene symbols were updated according to our dataset. Moreover, we used these gene lists to show the correspondence between cluster and cluster marker genes. To do this, we created chord diagrams using the R package “circlize”, adapted according to Allen et al., 2020.

### Immunofluorescence

Larval brains were dissected and fixed in 4% paraformaldehyde for 30 min at room temperature; afterwards were washed 3 times with PBS with 0.1% Triton X-100 (PBT). Fixed brains were incubated for 20 min in PBS with 0.5% Triton X-100 and 1% Normal Goat Serum (blocking solution) and incubated with the primary antibodies over-night at 4°C. Next day, brains were washed 3 times, blocked 20 min and incubated with the secondary antibodies for 2h at room-temperature. Afterwards, brains were washed 3 times, incubated 10 min in PBS, mounted in aqua/polymount (Polysciences, Inc) and imaged.

Immunofluorescent images were acquired on a LSM880 (Carl Zeiss GmbH). Adobe Photoshop (Adobe) and/or Fiji were used to adjust brightness and contrast; figure panels were prepared in Illustrator (Adobe).

## QUANTIFICATION AND STATISTICAL ANALYSIS

### TapeStation

Quantification and quality of cDNA and respective libraries generated for 10x Genomics Data was assessed with TapeStation following the standard protocol available at https://www.agilent.com/cs/library/usermanuals/public/ScreenTape_HSD5000_QG.pdf (Agilent Technologies).

### Differential Expression between Clusters

Differential expression analysis was performed within TFs/DNAB only. We used Seurat to identify the specific markers for each cluster. For that, we used the Receiver Operating Curve (ROC) test to find the differentially expressed genes between clusters. Within that analysis, we selected an AUC (Area Under the ROC Curve) > 0.5, to assure that the only hits were from genes with predictive values to classify that cluster. Moreover, to assure that none of the hits is a scarcely expressed gene, we only considered genes expressed in at least 25% of the cells of either one of the groups compared. Furthermore, we established that the average log fold change between the two populations being compared should be higher than 0.4 (Figures 2C,D and **3,A,C,D**), except for the case presented next. For neuronal cluster comparison (**Figure S3A**), the average log fold change was altered to 0.25; moreover, the dot plots for the top cluster markers of these analysis only showed genes with a pct.1 > 0.5 and pct.2 < 0.2.

GO term enrichment analysis was performed in PANTHER with the statistical overrepresentation test for biological process.

The Venn Diagram (Figure 3E) was generated using the online tool molbiotools. By selecting the Multiple List Comparator we were able to compare the overlapping TFs/DNAB genes between the lists comparing type I NBs *vs.* type I GMCs and type II NBs *vs*. dINPs.

### Analysis of type II lineage morphology

Quantification of abnormal morphologies of type II lineages (control and TfIIFα RNAi) was performed manually in Fiji. Lineages were characterized regarding the existence of type II NBs and whether they were individualized from each other or mixed at the (d)INP/GMC level (Figure 3E). Statistical analysis was in GraphPad Prism version 6.01 (GraphPad Software, Inc.; La Jolla, CA, USA); statistical differences were determined using two-tailed paired Student’s *t* test. Data is presented as mean ± standard deviation (SD). p values < 0.05 were considered statistically significant.

### Quantification of cells expressing a gene (mRNA)

To identify the number of neurons expressing genes necessary for neurotransmitter biosynthesis (VGlut, VAChT, Gad1, Vmat), we determined the number of cells with more than 0 counts for each specific gene or combination of genes. Percentages were calculated in relation to the total number of neurons in the dataset.

## ACKNOWLEDGEMENTS

We are in debt with C. Desplan for the encouragement and the opportunity to perform the scRNA-Seq experiments. We thank F. Simon for his help during the scRNA-Seq protocol. We also thank C. Desplan and his group for experimental help and discussion regarding this study. We thank J. Knoblich and F. Bonnais for fly lines and antibodies. We thank A.M. Venda for providing IFs images for E(spl)-mγ-HLH expression in imGMCs. We thank all members of the C.C. Homem Lab and F. Pinto-Teixeira for helpful discussion. We also thank C. Mendes and R. Teodoro for revising the manuscript. We also thank CEDOC’s Microscopy and Fly facilities for technical support.

Stocks obtained from the Bloomington *Drosophila* Stock Center (NIH P40OD018537) were used in this study. The monoclonal antibodies were obtained from the Developmental Studies Hybridoma Bank, created by the NICHD of the NIH and maintained at The University of Iowa, Department of Biology, Iowa City, IA 52242. CEDOC’s Fly facility was funded by CONGENTO LISBOA-01-0145-FEDER-022170. This project has received funding from the European Research Council (ERC) under the European Union’s Horizon 2020 research and innovation programme (H2020-ERC-2017-STG-GA 759853-StemCellHabitat); by Wellcome Trust and Howard Hughes Medical Institute (HHMI-208581/Z/17/Z-Metabolic Reg SC fate); EMBO Installation grant (H2020-EMBO-3311/2017/G2017); by Fundação para a Ciência e Tecnologia (IF/01265/2014/CP1252/CT0004 and PD/BD/114253/2016 to GSM). N.K. was supported by the National Eye Institute (K99 EY029356-01). This work was also supported by an NYU Abu Dhabi Research Institute grant to the NYUAD Center for Genomics and Systems Biology (ADHPG-CGSB).

## CONFLICT OF INTEREST

The authors declare that they have no conflict of interest.

## AUTHOR CONTRIBUTIONS

G.S.M. and C.C.H designed the project. G.S.M. performed the genetic experiments and G.S.M. and C.C.H. analyzed the data. G.S.M and N.K. acquired the scRNA-Seq data and G.S.M. and J.T.R. performed bioinformatic analysis using Seurat, G.S.M. performed the analysis using Monocle and P.B. performed the analysis using Velocyto. G.S.M. and C.C.H. wrote the manuscript.

## Notes

### Competing Interest Statement

The authors have declared no competing interest.

