## Supplementary Figures for "Fate transitions in *Drosophila* neural lineages: a single cell road map to mature neurons"

**Supplementary Figure legends**

**Figure S1. Quality control analysis of transcriptome and cluster analyses. Related to Figure 1.**

(A) Quality control metrics showing UMI counts, feature counts and mitochondrial percentage in the initial dataset (before filtering).

(B) UMAP plot showing the uniform distribution of both samples used in the analysis.

(C) Feature plot with the feature counts for each cluster.

(D) Dot plot showing the expression of NB and neuronal specific markers in each cluster.

**Figure S2. Dynamics of differentiation in type I and type II neural lineages. Related to Figure 4.**

(A) Unspliced and spliced fractions identified for each sample. By analysing the balance between nascent (unspliced) mRNAs and mature (spliced) mRNAs in single cells, the RNA velocity method estimates an RNA velocity vector for each cell. This allows to predict the future state of every cell in a scale of hours.

(B) Velocity field projected in the UMAP plot; arrows indicate the future state of a cell. The analysis of RNA velocities nicely predicts the direction of lineage differentiation from the less differentiated NBs to the more differentiated neurons.

(C) Cell trajectory for type I NB lineages; cell ordering and trajectory defined based on pseudotime analysis using monocle; colored by cluster.

**Figure S3. Neuronal cluster marker profiles reveal diversity in neuronal fate identities. Related to Figure 5.**

(A) Top differentially expressed (DE) markers per neuronal cluster; pct.1 > 0.7 and pct.2 < 0.2.

(B) Number of neurons expressing neurotransmitter associated genes (*VGlut*, *Gad1*, *Vmat*, *VAChT*) independently or simultaneously (counts>0); the respective percentages within the total neuronal population are indicated in bold.

**Figure S4. ChAT expression in neurons with different ages. Related to Figure 6.**

(A) Schematic representations of 48h-clone induced in neural lineages. Clone induced at 87h after larval hatching (ALH) and analysed at approximately 24h after puparium formation (APF); (A') Immunofluorescence images, two Z-slices of same clone displayed. ChAT (purple), GFP (green); Scale bars, 20 μm.

**Figure S1. Quality control analysis of transcriptome and cluster analyses.**

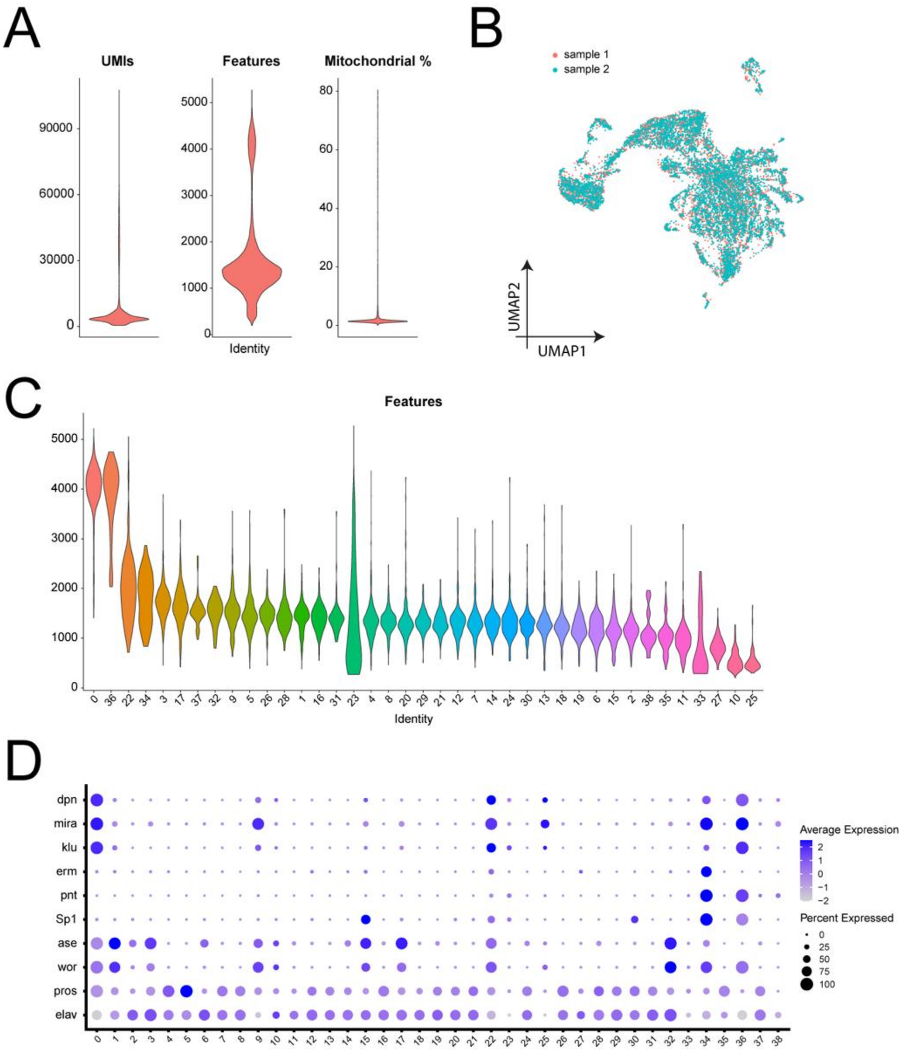

**Figure S2. Pseudotime analysis with Monocle and Velocyto.**

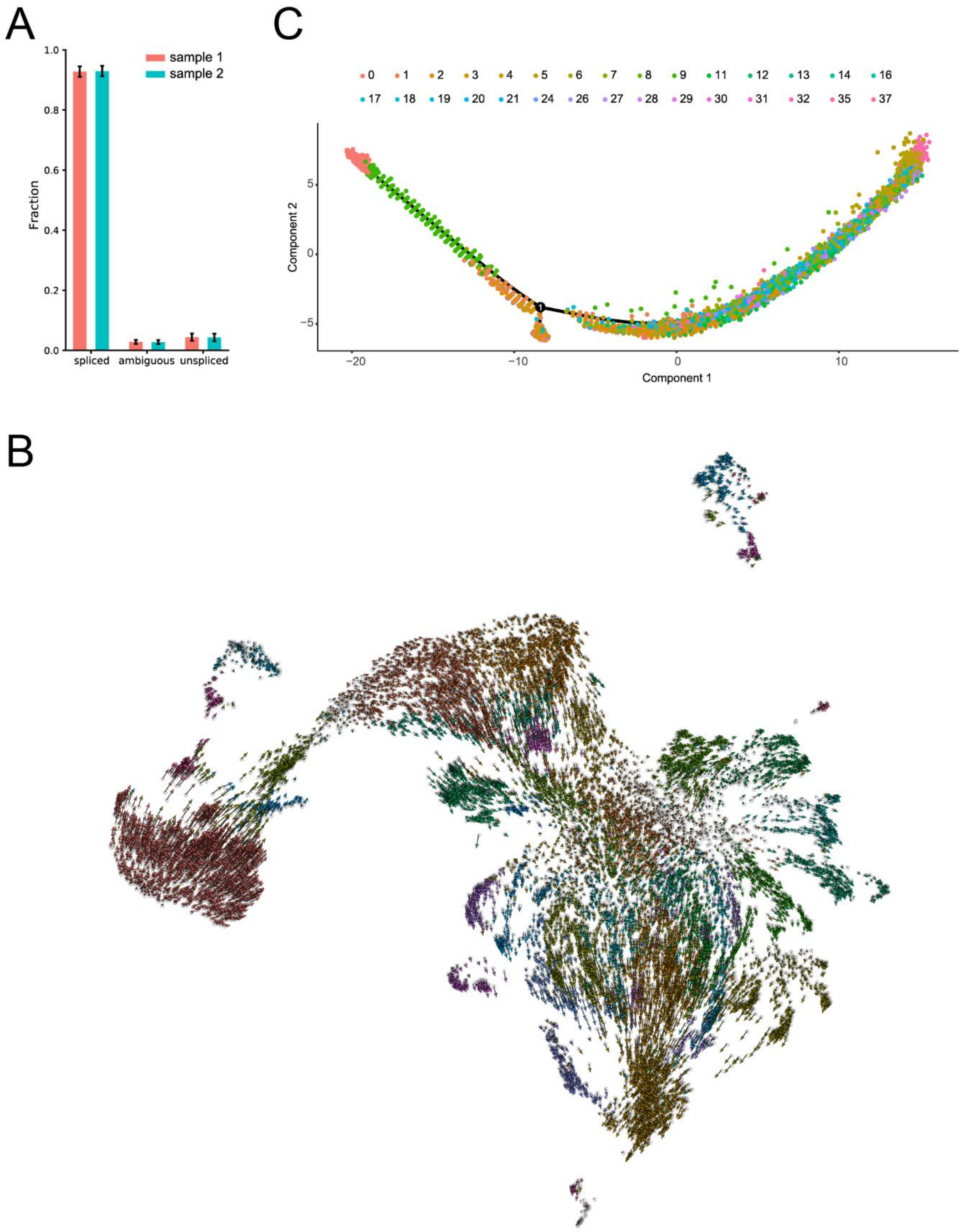

**Figure S3. Neuronal cluster marker profiles reveal diversity in neuronal fate identities.**

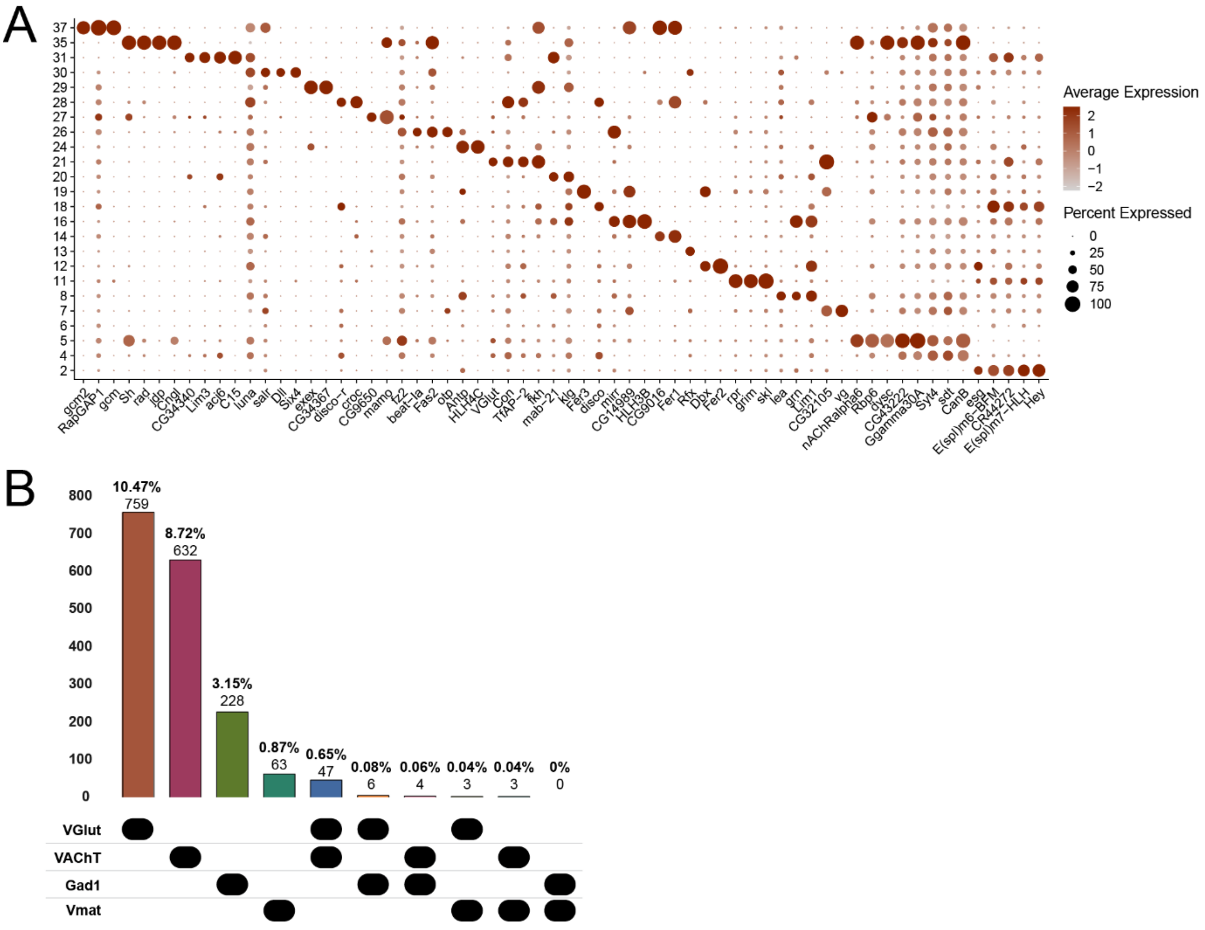

**Figure S4. ChAT expression in neurons with different ages.**

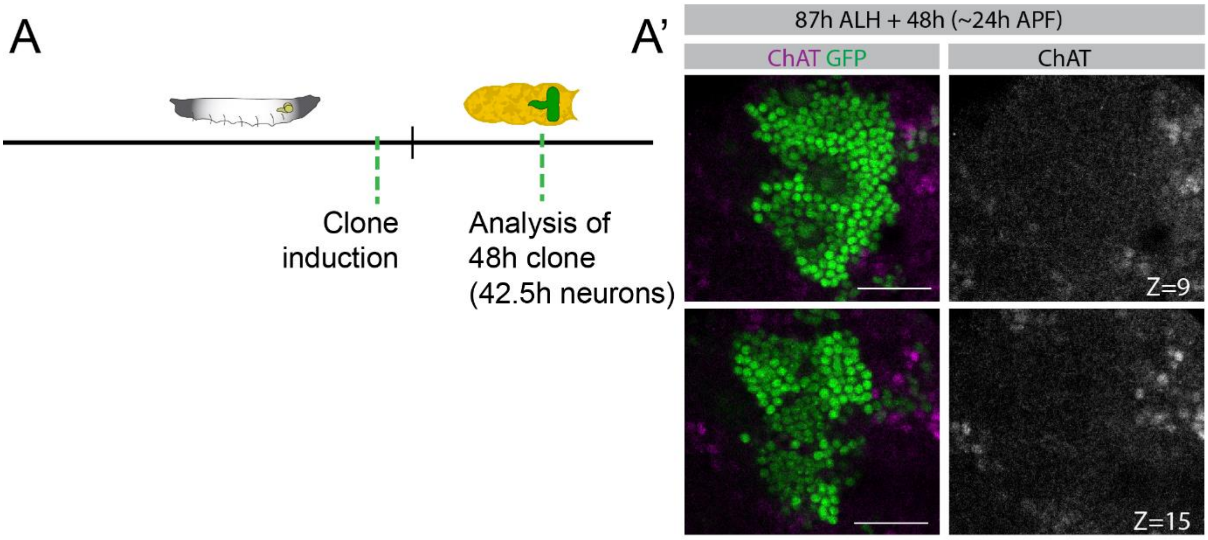
